# Diverse infections transcriptionally reprogram the intestinal epithelium and epithelial-immune cell interactions

**DOI:** 10.64898/2025.12.22.695505

**Authors:** Andrew Hart, Maria Merolle, Christian Howard, Breanne E. Haskins, Ian S. Cohn, Suhas Bobba, Rui Xiao, Yi Yang, Ken Cadwell, Junjie Ma, Hiroshi Yano, Xiaoxiao Hou, Bethan A. Wallbank, Daniel Cutillo, The MIST Consortium, Ivaylo I. Ivanov, Boris Striepen, Sunny Shin, Igor E. Brodsky, David Artis, Christopher A. Hunter, Daniel P. Beiting

## Abstract

The distal small intestine plays vital roles in host physiology by regulating nutrient and fluid homeostasis. Despite being impacted in Crohn’s disease and a major target for a range of infections, we know relatively little about the complexity of cellular responses and cell-cell communication in the ileum during infection. Single cell and spatial transcriptomics have emerged as powerful technologies to study tissue heterogeneity in the gut, but these tools have focused on the large intestine, in part due to the accessibility of this tissue for biopsies and its importance in cancer. Here we present GutPath, an atlas of over 500,000 single cells with RNA and protein expression profiles for 91 cell states in the ileum across diverse infectious archetypes. We show that GutPath accurately captures established immune responses to infection while revealing pathogen-specific responses in enterocytes. To highlight the discovery potential of this atlas, we identify a novel enterocyte cell state present during *Yersinia pseudotuberculosis* infection that is spatially linked to bacterial load and tissue pathology. GutPath establishes a much-needed resource for the immunology community that will accelerate the study of the transcriptional diversity of cellular landscapes in the small intestine.

## Introduction

The small intestine (SI) – organized into the duodenum, jejunum, and ileum – constitutes a massive surface area tasked with carrying out essential digestive functions while simultaneously contending with a battery of immunological challenges that include maintaining tolerance to food-derived antigens, regulating responses to commensals, and sensing and restricting infections^1–3^. To meet these physiologic and immune demands, the SI has evolved complex cellular environments with dozens of different cell types and unique metabolic and immune niches^4–6^. The ileum is distinct from other parts of the gut in that it plays a crucial role in nutrient and fluid balance as the primary site for absorption of vitamin B12 and bile acids, as well as being highly efficient at absorption of fat-soluble vitamins (A, D, E, and K), water, sodium, and chloride^7,8^. The epithelium of the ileum regulates nutrient transport and is a physical and chemical barrier to infection. This barrier is formed by a single layer of epithelial cells comprised of absorptive enterocytes, goblet cells and their secreted mucins, Paneth cells (highly enriched in the ileum) producing antimicrobial peptides, tuft cells, and other specialized populations that collectively maintain gut homeostasis. Epithelial cell diversity in the gut varies between the SI and the colon, between different anatomical regions of the SI, and even within microanatomical structures such as villi^4,9,8^. Intestinal stem cells located in the villus crypts give rise to nascent absorptive enterocytes that produce antimicrobial peptides and have immune functions. As these cells mature and advance toward the villus tips, tightly regulated transcriptional programs alter their functional capacity to perform nutrient transport^4,10^. Interspersed with and underlying the epithelial barrier in the lamina propria (LP) are diverse immune cells that aid in immune tolerance and pathogen defense^6^. Finally, stromal populations and the enteric nervous system are essential for tissue architecture, cell-cell communication, and gut physiology and further contribute to the cellular heterogeneity of the SI.

Because of the vital role played by the ileum in nutrient and fluid balance, inflammatory and infectious diseases that affect this site can lead to major disruptions to host metabolism and physiology. For example, in patients with Crohn’s disease, the ileum is a commonly affected part of the gastrointestinal (GI) tract and as many as a third of all patients present with exclusive ileal inflammation^11^, associated with malabsorption of vitamin B12 and bile acids^12,13^. The ileum is also a common target for pathogens that can have severe ramifications that range from diarrhea and growth delays to death^3,14,15^. Cryptosporidiosis, caused by the apicomplexan parasite, *Cryptosporidium* spp., affects the small intestine and is a leading cause of diarrheal mortality and growth stunting in children, globally^14^. Bacterial pathogens, including *Salmonella*, *Shigella*, *Campylobacter*, *Listeria, Yersinia* and some pathogenic *E. coli*, gain access through Peyer’s patches that are abundant in the ileum. Likewise, many helminth parasites establish infection in the ileum and cause significant morbidity worldwide^15^. Pathogens that infect the ileum vary in physical size, immunogenic molecular patterns, cellular tropism, and pathogenicity. Accordingly, the immune response elicited by each pathogen is highly tailored to optimize defense and ensure survival. Despite the complex and multifaceted roles of the ileum in host biology and defense and the breadth of infection-driven transcriptional responses, relatively little effort has been devoted to single cell analysis of this gut region. Studies focused on infections of the small intestine including *Salmonella*, Astrovirus, and *Cryptosporidium* have informed our understanding of how tissue specific immunity is performed and regulated in the gut but are often limited to one or a few cell types and cannot easily be extrapolated or compared to other infection systems^4,16–18^. Thus, there is an unmet need for large-scale single cell studies that interrogate the distal small intestine across a wide range of pathogenic phyla to capture how distinct organisms alter cellular heterogeneity and how pathogen-specific immunity is coordinated through transcriptional regulation.

The development of single cell technologies designed to capture RNA (scRNA-seq), protein, and epigenetic data have expanded our understanding of mucosal physiology and immunology. Recent single cell transcriptional studies of the gut have led to the discovery of previously unidentified cell states^4,7,9,10,19–23^, including novel enteroendocrine cell populations and inflammation-associated changes to the colonic intestinal stem cell pool during colitis^4,22^. To date, intestinal scRNA-seq studies often deeply profile specific cell populations^10,24–26^ or catalogue the steady state and developmental conditions of humans^7,9,20,23^ or mice^4,19^. Intense effort has focused on the large intestine where ulcerative colitis and colon cancer drive research interests and biopsies are readily available for human subject research^22,25,27–31^, and where extensive work has been devoted to develop mouse models of inflammation^32,33^. Although these data capture GI biology at steady-state and during various diseases, infection-induced immune processes are underrepresented^20,19,22,34^. Consequently, there is a need to better understand the unique transcriptional and immune responses accompanying infection with distinct classes of pathogens.

To address this critical knowledge gap, we created GutPath (available at GutPath.org), a single cell RNA and protein (CITE-seq) atlas constructed from the ileum across a range of viral, bacterial, fungal, protozoan, and helminth mouse models of infection and colonization. This resource contains more than 500,000 cells from the ileum and ileal-draining mesenteric lymph node (MLN). 91 annotated cell states in the ileum and 49 cell states in the MLN were organized into multiple layers of reference annotation from broad cell lineages to refined and descriptive cell states. We demonstrate that GutPath accurately captures acute infection biology for six model pathogens that impact the distal small intestine. Furthermore, we identify pathogen-specific responses including unique enterocyte transcriptional responses and cell-cell signaling programs that arise during *Nippostrongylus brasiliensis* and *Yersinia pseudotuberculosis* infections. Finally, using spatially resolved single cell transcriptomics we illustrate that unique enterocyte responses develop locally, regionally and tissue-wide during infection.

## Results

### Creating a single-cell atlas of the small intestinal response to diverse infections

GutPath was constructed to fill critical gaps in our understanding of intestinal biology and to create a resource that captures cellular heterogeneity during homeostasis and innate immune responses to mucosal infection. We focused on microbes that infect or colonize the small intestine while sampling a diverse range of species from distinct phyla, including: Segmented filamentous bacteria (SFB, *Candidatus savagella* spp: gram-positive commensal bacterium), *Yersinia pseudotuberculosis* (gram-negative bacterium), *Nippostrongylus brasiliensis* (multicellular eukaryotic parasitic helminth), *Candida albicans* (fungus), *Cryptosporidium parvum* (unicellular apicomplexan parasite), and murine norovirus strain CR6 (MNV: positive-strand ssRNA Caliciviridae virus) **(Figure 1)**. All infections or colonizations were orally administered, apart from subcutaneously injected *Nippostrongylus* L3 larvae, and tissues were collected during the first week of infection **(Figure 1)**. Single cell RNA and ∼100 surface proteins were simultaneously profiled using Cellular Indexing of Transcriptomes and Epitomes (CITE-seq) of the MLN, the small intestine lamina propria (LP), and the small intestine epithelial cell (IEC) layer for at least two replicate mice per condition. Additional duodenal LP and IEC samples were sequenced from *Nippostrongylus-*infected mice to capture the site where worm burden is highest^35^.

**Figure 1:**
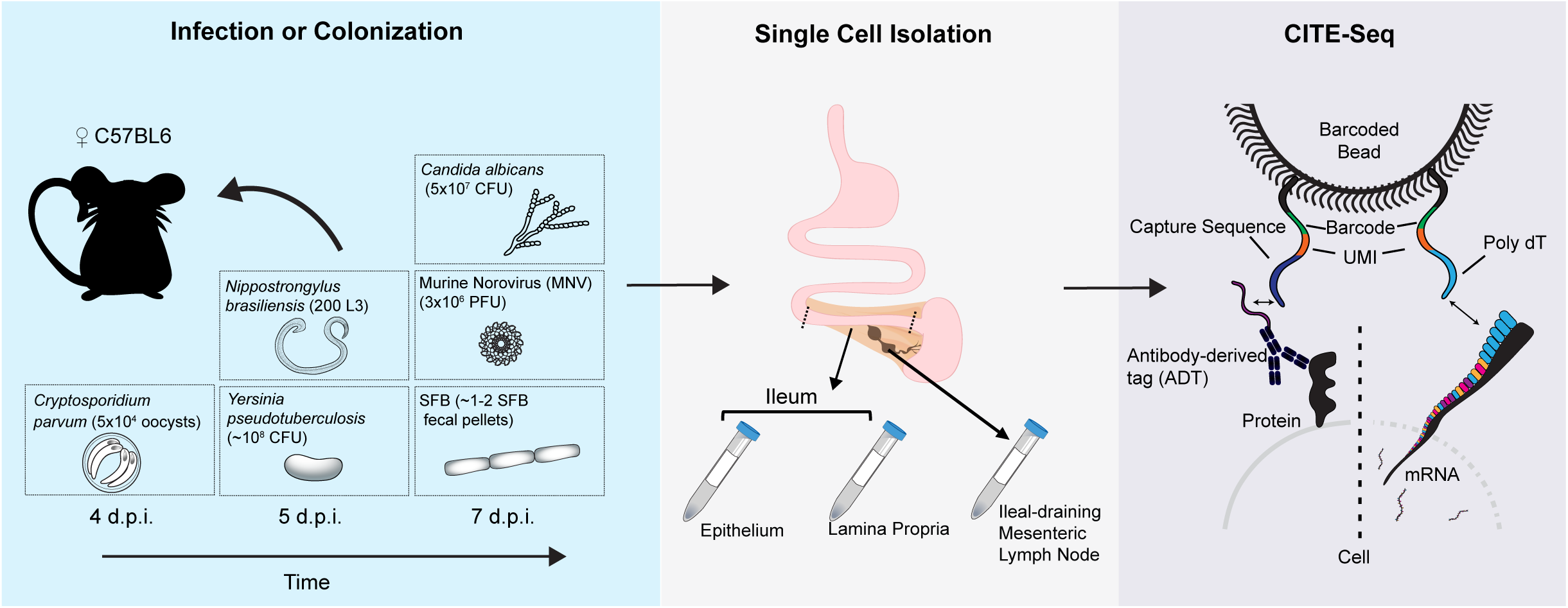
Generating GutPath. Experimental model depicting three major phases of GutPath sample generation: infection, tissue and single cell isolation, and library sequencing. C57BL6 mice were infected/colonized orally or subcutaneously with one of 6 infectious or colonizing organisms. Tissue samples were taken between four- and seven-days post infection from the distal third of the small intestine and the ileal draining mesenteric lymph node. Separate single cell samples and libraries were prepared for the intestinal IEC and LP compartments. Sequencing libraries of antibody-derived tag expression and mRNA expression were prepared according to 10x Genomics protocols and were sequenced.

Across MLN, IEC, and LP fractions, mean reads per cell exceeded 30,000 for mRNA and 8,000 for Antibody-Derived Tags (ADT) **(Figure S1A and B, respectively, dotted lines)**. After quality control analysis **(Figure S1C; Methods)** we retained 505,956 high quality cells that were subsequently separated by tissue type for downstream analyses – 184,973 and 320,983 cells from the MLN and ileum, respectively. These cells contained on average 5,867 UMI transcripts/cell and 2,258 genes/cell in the MLN; and 6,398 transcripts and 2,186 genes/cell in the ileum **(Figure S1D, dotted line; and Figure S2A)**. At least 40,000 and up to 100,000 cells were collected per condition across tissues **(Figure S2B)**. To visualize these cells in a two-dimensional space, mRNA and ADT data were integrated independently and combined through weighted nearest neighbor analysis **(Figure 2A and 2B)**. Cell types were classified using automated methods complemented with manual curation and validated by gene and protein expression profiles **(Figure S2C-E)**. As expected, the MLN was dominated by B cells, T cells, and ILC populations (collectively, 95% of MLN cells) **(Figure 2C and Figure S2F).** Importantly, while epithelial cells comprised 80% of the IEC fraction **(Figure 2D and Figure S2G)**, further digestion of the SI to recover the LP fraction yielded additional epithelial populations, particularly Paneth and stem cells, and transit amplifying cells **(Figure 2D and Figure S2G)**. Compared to the IEC fraction, LP was enriched in mesenchymal, endothelial, cells of the enteric nervous system, and B cells **(Figure S2G)**. We also observed infection-specific alterations in cellular composition of the LP fraction, with *Yersinia* inducing expansion of neutrophils, while *Nippostrongylus* infection led to expansion of Tuft cells and ILCs **(Figure S2G)**.

**Figure 2.**
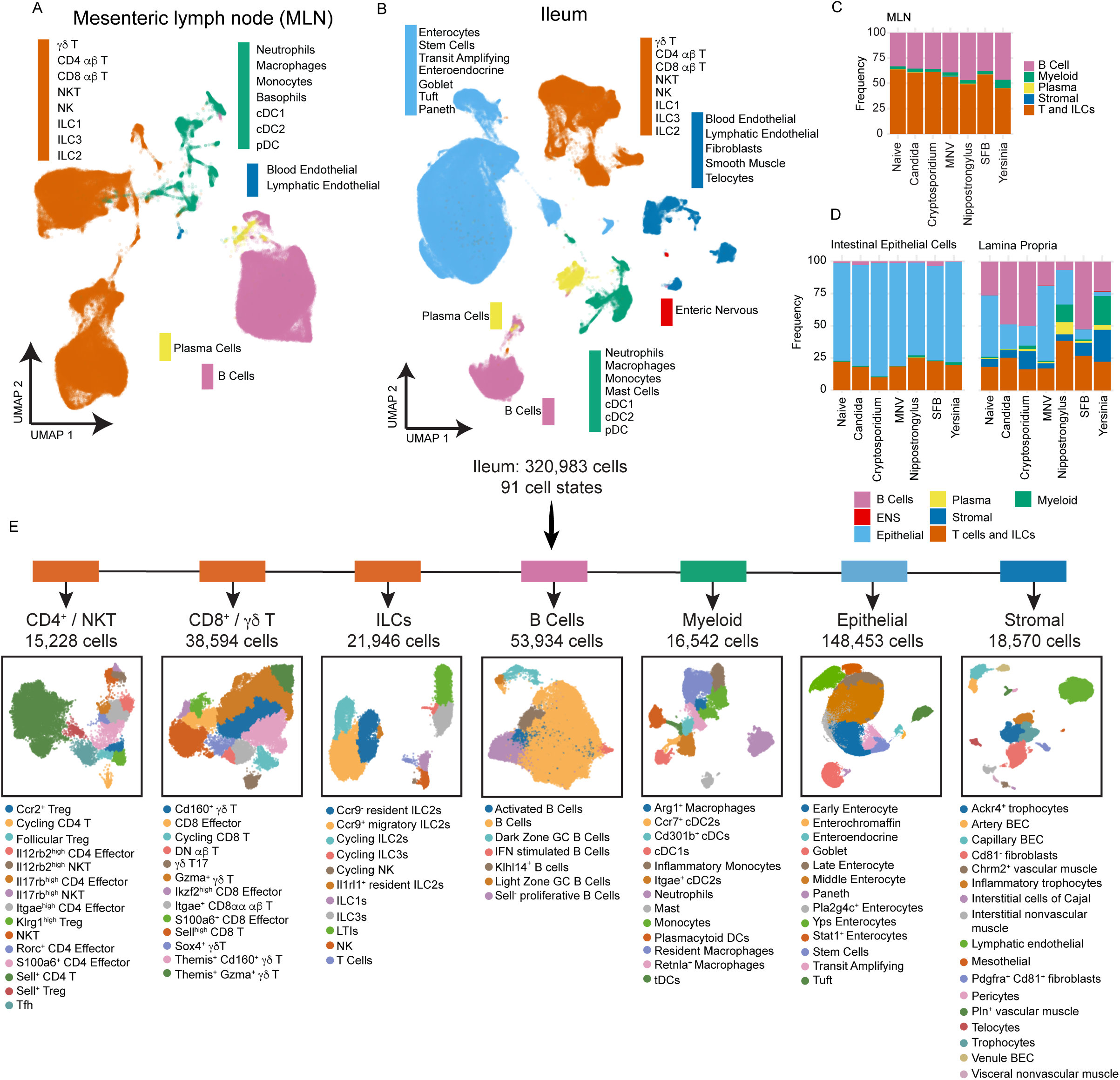
: GutPath is a robust measure of mucosal responses to infection. A) CITE-seq UMAP dimensional reduction depicting major cell lineages (color) and labeled annotated cell types in the MLN. B) CITE-seq UMAP dimensional reduction depicting major cell lineages (color) and labeled cell type annotations in the Ileum across all infections and conditions. C) Frequencies of major cell lineages in the MLN for each condition. D) Frequencies of major cell lineages (color) in the IEC (left) and LP (right) data across infections/colonizations. E) Schematic UMAP diagrams of ileum subsets that were segregated, re-clustered, and subjected to refined cell type annotation. Cell numbers and refined cell type annotations of each subset are provided. Major cell lineages to which each subset belongs are indicated (top) by color as appearing in B. Plasma cells and enteric nervous system lineages were not further subset and are not shown. Biological n: Naive = 2, Candida = 2*, Cryptosporidium = 3, MNV = 2, Nippostrongylus = 2**, SFB = 2, Yersinia = 3. *Candida infection includes 1 IEC replicate. **Nippostrongylus biological replicates include both ileal and duodenal-derived lamina propria and epithelial layer samples.

The size and cellular complexity of the small intestine makes cell type annotation critical to fully capture tissue heterogeneity, yet even with the advent of improved automated methods high-resolution cell annotation remains a major challenge^4,9,22,36^. We utilized existing resources and extensive manual annotation to generate highly resolved cell labels for our atlas that defined 49 and 91 separate cell states in the MLN **(Figure S3A)** and ileum **(Figure 2E)**, respectively. At the most resolved level of cell state description, we can delineate early, middle, and late stages of enterocyte differentiation^10^, as well as infection-specific enterocyte states in mice infected with *Yersinia* (hereafter, referred to as ‘Yps enterocytes’) or *Nippostrongylus* (*Pla2g4c*+ enterocytes) **(Figure 2E)**. In the *Yersinia-*infected intestine we also observed an inflammatory population of trophocytes (*Pdgfra*^lo^ and expressing *Il6*, *Ccl2*, and *Cxcl1*), a stromal cell type that supports intestinal stem cells (ISCs) and regulates their differentiation^37^. Our atlas also captured several rare cell types including telocytes (*Pdgfra*^high^), a population of stromal cells that also coordinate ISC Wnt signaling and differentiation^38^; pericytes (*Cspg4*^+^ and *Pdgfrb*^+^) that stabilize vessels and support blood flow; enterochromaffin cells (*Lmx1a*^+^), a subset of enteroendocrine cells that produce serotonin; and interstitial cells of Cajal (*Kit*^+^), which are the motility “pacemakers” of the gut **(Figure 2E)**^39,40^. The granular annotation of cell states in GutPath constitutes a valuable resource that can be used for label transfer in mucosal studies, thus accelerating the analysis of single cell data in the field. To make these cellular descriptions accessible to researchers, complete marker profiles were generated for coarse cell lineages, intermediate cell types, and the most refined cell state annotations **(Supplementary Table 1)** using a differential expression approach^41^. Additionally, we utilized a random forest-based machine learning approach to create highly condensed, necessary and sufficient, marker profiles for each cell state and level of annotation **(Supplementary Table 2)**^42^.

### GutPath captures immune hallmarks of model infections

To test whether GutPath faithfully recapitulates pathogen-specific innate immune responses, we utilized differential abundance testing of cell ‘neighborhoods’ (defined by k-nearest neighbor) to look for both differential abundance of cell populations and the transcriptional heterogeneity in those populations relative to naive samples. Differentially abundant neighborhoods containing tuft cells and goblet cells were enriched in *Nippostrongylus* infection **(Figure 3A)**, but not other infections, consistent with the tuft and goblet cell hyperplasia that is a feature of helminth infections **(Figure S4A-B)**^43^. Rapid recruitment of monocytes, neutrophils, and other myeloid effector cells is a dominant feature of many bacterial infection models including *Yersinia pseudotuberculosis*^44^ and we observed differentially abundant neighborhoods and increased frequencies of myeloid populations in the *Yersinia*-infected samples **(Figure 3B and Figure S4C-D)**. Monocyte neighborhoods were also differentially abundant in SFB colonization, another bacterial model, and *Nippostrongylus* infection resulted in neighborhoods of moderately increased neutrophil and monocyte abundance.

**Figure 3:**
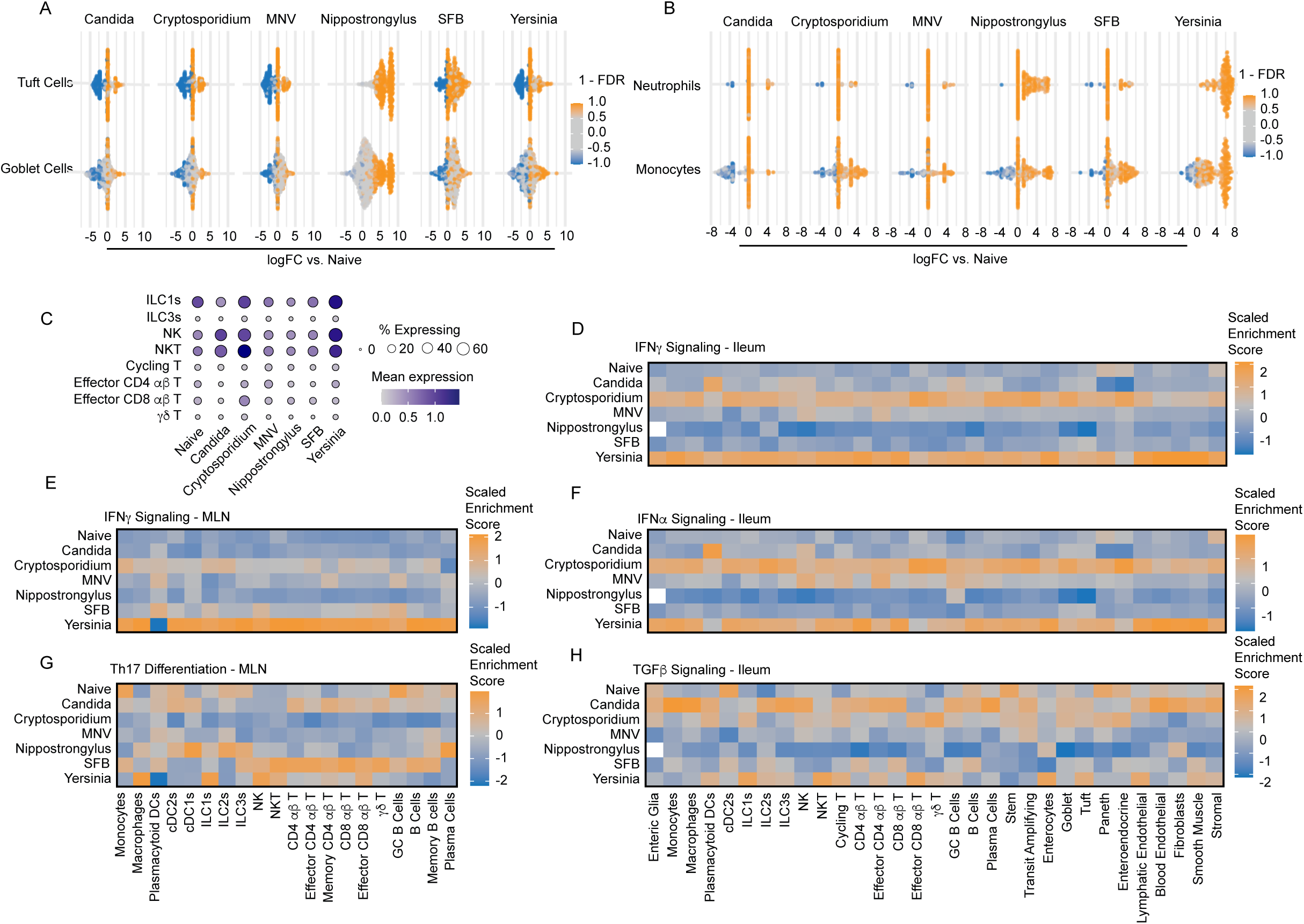
Infection paradigms are faithfully recapitulated in GutPath. A) Differential abundance analysis of defined cell neighborhoods within tuft cells and goblet cells of the ileum. Comparisons between each infection and naive samples are shown with significantly altered neighborhoods indicated by color (orange = increased in infection/colonization, blue = decreased in infection/colonization). B) Differential abundance neutrophil and monocyte neighborhoods in the ileum. Comparisons between each infection and naive samples are shown. Significantly altered neighborhoods are indicated by color (orange = increased in infection/colonization, blue = decreased in infection/colonization). C) Frequency of cells expressing *Ifng* (bubble size) and the expression values of Ifng (color scale) across cell types and in each condition within the ileum. D) Scaled UCell signature scores of the Hallmark IFN-γ signaling gene signature across ileum cell types in each infection. E) Scaled UCell signature scores of the Hallmark IFN-γ signaling gene signature across MLN cell types in each infection. F) Scaled UCell signature scores of the Hallmark IFN-ɑ signaling gene signature across ileum cell types in each infection. G) Scaled UCell signature scores of the GO:BP Th17 differentiation gene signature across MLN cell types in each infection. H) Scaled UCell signature scores of the Hallmark TGF-β signaling gene signature across ileum cell types in each infection. Biological n: Naive = 2, Candida = 2*, Cryptosporidium = 3, MNV = 2, Nippostrongylus = 2**, SFB = 2, Yersinia = 3. *Candida infection includes 1 IEC replicate.

*Cryptosporidium* infection elicits IFN-γ signaling and IFN-γ-dependent immunity, and we observed increased *Ifng* expression in NK and NKT cells at this early time point in the *Cryptosporidium* and *Yersinia*-infected samples (**Figure 3C)**. We also tested whether increased expression of cytokine resulted in increased IFN-γ signaling as measured by enrichment scores of the Hallmark IFN-γ signaling gene set. In the ileum, there was activation of the IFN-γ signaling in cells of both the *Cryptosporidium* and *Yersinia*-infected mice **(Figure 3D)**^45^. Modest enrichment was observed in the MLN of *Cryptosporidium* infected samples while enrichment of IFN-γ signaling was robust in MLN cells from *Yersinia-*infected mice, likely due to the systemic nature of this infection **(Figure 3E)**^46^. Next, persistent CR6 strain of MNV is sensitive to STAT1-driven immunity despite producing a less robust type 1 IFN response compared to other MNV strains^47,48^. However, in NK cells and CD4^+^ T cell populations, there was an enrichment of the type 1 IFN signaling gene set during MNV infection **(Figure 3F)**.

*Candida* colonization is complex and can be pathologic or commensal in mice, in part through interactions with the microbiome^49^. Negative effects of *Candida* colonization have been linked to local induction of hypoxia in the intestine^50^, and we observed moderate enrichment of hypoxia-related gene signatures in cells from C*andida* colonized mice **(Figure S4E)**. *Candida* is also known to promote the development of Th17 responses. We did not observe robust Th17 responses in *Candida* in the ileum at 7dpi. However, TGF-β signaling – a cytokine which drives Th17 differentiation and is systemically increased during *Candida colonization*^51,52^ – was enriched in cells from *Candida-colonized* mice **(Figure 3H)**. Additionally, in the MLN, *Candida* colonization led to relative enrichment of the Th17 differentiation gene set in T cell populations. SFB colonization, another Th17-inducing model, likewise showed elevated Th17 differentiation gene set enrichment in T cell populations compared to naive samples and other infection models **(Figure 3G)**^53^. Thus, these data show that GutPath accurately captured immunological archetypes of phylogenetically-diverse model infections.

### Infection alters enterocyte development and nutrient physiology

Given the clear induction of canonical immune pathways across the six infections **(Figure 3)**, we next performed unbiased model-based differential gene expression analysis to identify unique responses to infection in the SI, compared to naïve animals^54^. We hypothesized that many pathways associated with tissue damage or with general immune activation, including proliferation, might be conserved across different infections and cell types. Alternatively, since GutPath captures innate immune responses to phylogenetically diverse pathogens – and the immune system recognizes evolutionarily conserved molecular patterns – transcriptional responses to infection could be cell type- and pathogen-specific. The magnitude of transcriptional changes varied greatly across infections and between cell types. While MNV had the fewest total number of DEGs, *Candida* colonization showed the weakest induction of DEGs as measured by logFC **(Figure 4A and Figure S5A)**. Compared to MNV and *Candida*, *Cryptosporidium* and SFB both elicited more DEGs and more dramatic induction of these DEGs, while *Nippostrongylus* and *Yersinia* infection resulted in the most robust transcriptional response **(Figure 4A and Figure S5A)**. Across the infections, these responses mapped primarily to enterocytes and γδ T cells. However, when we restricted the analysis to genes more strongly dysregulated during infection (logFC ≥ 1.5), IECs including enterocytes, stem, transit amplifying, and goblet cells, as well as fibroblasts were identified as major drivers of the innate response across all infections, having the most DEGs on average compared to other cell types (Figure 4B)

**Figure 4:**
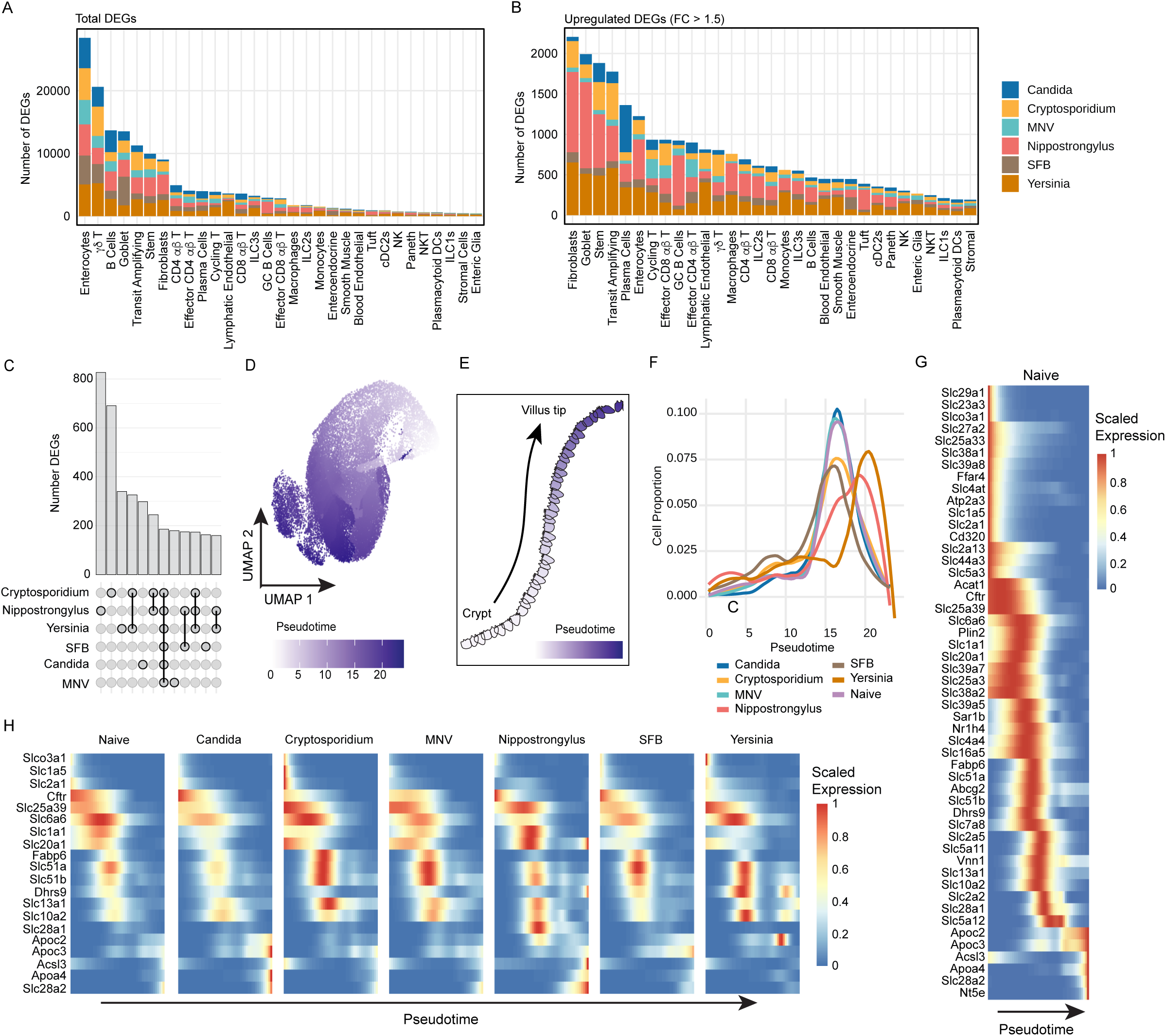
Enterocyte physiology is disrupted during infection. A) Cumulative MAST differential expression results for each cell type (bars) and infection (colors) in the ileum. DEG counts (FDR <0.05) are shown. B) Cumulative MAST differential expression results for each cell type (bars) and infection (colors) in the ileum. DEG counts (FDR <0.05 and FC > 1.5) are shown. C) Upset plot co-occurrence of upregulated DEGs from stem cells, transit amplifying cells, and enterocytes across all infections. Overlapping gene counts (top, bars) and infection overlap key (bottom) for the 12 most numerous combinations are shown. D) UMAP reduction indicating pseudotime progression (color hue) of subset and reclustered stem cells, transit amplifying cells, and enterocyte populations from all conditions. E) Small intestine villus diagram of pseudotime trajectory (color hue) representing stem to enterocyte differentiation. F) Proportional abundance of enterocyte lineage cells (stem cells, transit amplifying cells, and enterocytes) across pseudotime for each infection (color). G) Pseudotime-scaled expression of selected solute transporter and nutrient-associated genes identified as significantly DE along pseudotime among naive stem cells, transit amplifying cells, and enterocytes. H) Pseudotime-scaled expression of solute transporter and nutrient-associated target genes for each condition. Target genes differentially expressed (adj. p <0.05) in at least one infection relative to naive are shown. Biological n: Naive = 2, Candida = 2*, Cryptosporidium = 3, MNV = 2, Nippostrongylus = 2, SFB = 2, Yersinia = 3. *Candida infection includes 1 IEC replicate. BH multiple comparison correction was applied to MAST DE analysis and FDR <0.05 was applied.

Absorptive enterocytes provide the first cellular barrier to SI infection but are also critical regulators of host metabolism and have unique physiology that allows them to transport lipids, amino acids, and nucleotides from the lumen of the intestine into the tissue^10^. The large number of IECs in GutPath together with our finding that enterocytes, stem cells, and transit amplifying cells elicit robust transcriptional responses to infection prompted us to further investigate pathogen-specific and conserved responses to infection in these populations. Strikingly, we observed that DEGs identified in stem cells, transit amplifying cells, and enterocytes showed little overlap between infections and suggested substantial pathogen-specific regulation **(Figure 4C)**. We reasoned that pathogen-specific responses in IECs might be explained by alterations of enterocyte differentiation. Stem cells in the base of the crypts progressively differentiate from stem cell, to transit amplifying cells, and ultimately into immature enterocytes which continue to mature and alter their gene expression as they move to the tip of the villus^10^. Using pseudotime trajectory inference, we captured this maturation process **(Figure 4D-E** and **Figure S5B)**. To validate this trajectory among naive cells and across infections, we examined marker genes with well-established expression patterns during enterocyte differentiation, including *Lgr5* (stem cells), *Reg3b* (early enterocyte), *Slc5a1* (late enterocyte), and *Ada* (mature enterocytes at villus tip). All four genes showed the anticipated ‘timing’ of expression in naïve mice and across all infections combined **(Figure S5C)**^10^. Next, we evaluated the distribution of enterocytes along this pseudotime to gauge how each infection shapes enterocyte differentiation. Colonization with *Candida* and MNV infection did not shift enterocyte pseudotime, compared to Naïve mice **(Figure 4F)**. We observed a loss of mature enterocytes during infection with *Cryptosporidium*, a pathogen that exclusively infects enterocytes^16^, and a similar phenotype was observed in SFB colonization **(Figure 4F)**. In contrast, *Yersinia* and *Nippostrongylus* infections, which elicited the most DEGs in enterocytes **(Figure 4B)**, shifted pseudotime forward, indicating that these infections may drive enterocytes toward a more mature state or distinct transcriptional states.

Results highlighting enterocyte pseudotime regulation by infection prompted us to explore enterocyte gene expression along pseudotime in more detail. We employed tradeSeq^55^, a generalized additive model framework based on the negative binomial distribution, to analyze enterocyte gene expression along pseudotime during infection. We used naive samples as a baseline to define temporally regulated DEGs along the enterocyte pseudotime (n = 4,278). To probe the nutrient transport and metabolic regulatory capacity of cells along pseudotime, we cross-referenced the 4,278 enterocyte DEGs with a list of 134 metabolic and nutrient-regulating genes (**Supplementary Table 3**) including many solute carriers^56^. Solute carrier transporters are transmembrane proteins enriched on the apical and basal surfaces of enterocytes and their dysregulation leads to metabolic disease^10,57^. There is evidence that solute carriers are actively regulated during infection and participate in pathogenesis^58^, but their role in infection biology has not been explored across models. From our list of 134 solute carrier and metabolic regulator-genes, 51 were differentially expressed across pseudotime in naïve mice **(Figure 4G)**. Among these 51 genes, 20 (39%) were differentially expressed in at least one infection relative to naïve samples **(Figure 4H)** and generally followed one of two patterns. The first pattern is evident with genes such as the neurotransmitter transporter, *Slc28a2*, and the prostaglandin and bile acid transporter, *Slc6a6* which maintained the same ‘timing’ across enterocyte development as observed in naïve mice, but were downregulated by *Yersinia* and *Nippostrongylus* infections, respectively **(Figure 4H)**. A second pattern occurred when infection altered the timing of expression, rather than magnitude, which was observed with genes such as *Slc20a1*, a sodium-phosphate symporter that regulates absorption of phosphate from interstitial fluid; and *Slc25a39*, a mitochondrial protein responsible for uptake of glutathione **(Figure 4H)**. Together, these data provide a detailed ontological map of genes involved in enterocyte physiology and underscore the impact of infection on the timing and magnitude of their expression

### Enterocytes exhibit pathogen-specific responses during infection

The changes we observed in solute transporter expression prompted us to assess how metabolic pathways might be altered in enterocytes during infection. We utilized scCellfie to score distinct metabolic ‘tasks’ – a specific functional activity or purpose that a cell performs using metabolic pathways – across our cell populations and infection models^59^. This analysis showed that enterocytes from *Nippostrongylus*-infected mice exhibited the most dramatic alterations to metabolic tasks, particularly in pathways involved in lipid metabolism **(Figure 5A).** This included reduced synthesis of several secondary bile acids **(Figure 5B, left)** and increased degradation of multiple fatty acids, such as elaidate, linoleate, palmitate and arachidonate **(Figure 5B, right)**. Given the vital role played by enterocytes in regulating solutes and metabolism, we hypothesized that this phenotype would be restricted to the epithelial compartment. Indeed, the median difference in activity score for select fatty acid-related tasks between *Nippostrongylus* and naive samples across cell types was highest in enterocytes, irrespective of their maturity, and goblet cells **(Figure 5C)**. We used cell-cell communication analysis to identify the signaling events that might drive the metabolic dysregulation in *Nippostrongylus-*infected mice. Compared to enterocytes in naive mice, enterocytes in *Nippostrongylus-*infected mice are predicted to receive IL4 and IL13 signals **(Figure 5D, right, arrow; and Figure 5E)**, as expected of a T helper type 2 (Th2) dominant immune response. These Th2 cytokines drive tuft cell and goblet expansion from ISCs, but their effects on functional capacity and signaling in absorptive enterocytes, which express the appropriate receptors to respond, is not understood^60^. Enterocytes from *Nippostrongylus-*infected mice were also predicted to produce unique ligands for cell-cell interactions not found in naive mice including FGF family ligands (**Figure 5D, left, arrow**). FGF family and CX3C family ligands, specifically Fgf15 and Cx3cl1, are produced by enterocytes and were inferred to act on other cell populations **(Figure 5F).** While Cx3cl1 is notable for its chemoattractant capacity for mast cells, Fgf15, fibroblast growth factor 15, is a critical regulator of bile acids and cholesterol metabolism^61^. Taken together, these data suggest that infection with *Nippostrongylus* reprograms the metabolic activity of IECs in the ileum and that Th2 cytokines may have distinct effects on absorptive enterocytes that have yet to be explored.

**Figure 5:**
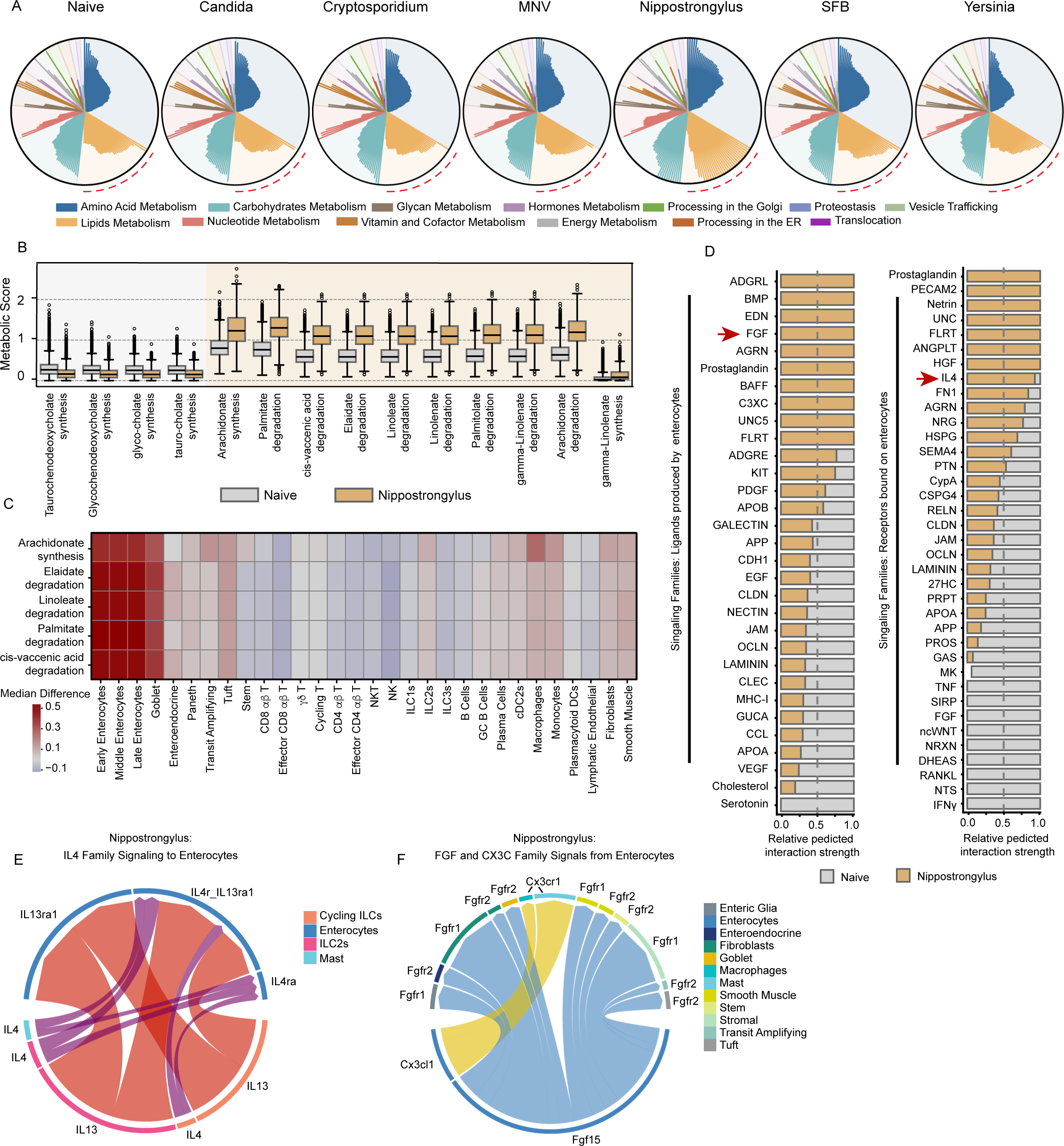
Nippostrongylus infection results in unique disruptions to enterocyte metabolic capacity. A) ScCellfie metabolic task scores for enterocytes from each infection. Radial plots depicting scaled activity scores across metabolic categories (colors) and individual metabolic tasks (bars). B) Significantly altered (adj. p <0.05, absolute Cohen’s threshold > 0.5, absolute logFC > 0.585 (∼1.5 FC)) lipid metabolism task score distributions for enterocytes of *Nippostrongylus*-infected (yellow) and Naive (grey) samples. C) Difference of medians heatmap showing metabolic task score activity change across cell types between *Nippostrongylus*-infected and Naive samples. D) Cell-cell communication signaling predictions for all signals produced by enterocytes (left) or received by enterocytes (right) of *Nippostrongylus*-infected mice relative to signals predicted in enterocytes of Naive mice. Significantly different predictions are shown (p <0.05) and FGF family and IL4 family signals are highlighted (red arrows). E) Chord diagram of IL4 family signals produced by various cell types (circumference bar colors) in *Nippostrongylus*-infected mice and received by IL4 family receptors on enterocytes. F) Chord diagram of FGF and CX3C family signals produced by enterocytes from *Nippostrongylus*-infected mice and received by various cell types (circumference bar colors). (Biological n: Naive = 2, Candida = 2*, Cryptosporidium = 3, MNV = 2, Nippostrongylus = 2, SFB = 2, Yersinia = 3. *Candida infection includes 1 IEC replicate.

### *Yersinia* infection stimulates a unique pro-inflammatory program in enterocytes

Given the dramatic impact of *Nippostrongylus* infection on enterocytes and the fact that both *Nippostrongylus* and *Yersinia* infection had the largest number of enterocyte-derived DEGs of all the infection models examined **(Figure 4A-B)**, we next questioned how *Yersinia* infection impacts enterocyte biology. Focusing on just the enterocyte cluster from GutPath, we first examined the extent to which each infection produced a similar clustering pattern, measured as a scaled imbalance^62^. This analysis showed that *Yersinia*, but not other infections resulted in an imbalance in the enterocyte clustering, suggesting that *Yersinia* may give rise to unique transcriptional states in enterocytes **(Figure 6A)**. To explore this further, we applied tradeSeq to determine differential expression along enterocyte pseudotime compared to naïve animals and found that *Yersinia* elicited more temporally regulated DEGs than all other infections combined (adj p. < 0.05 & logFC > 2) **(Figure 6B)**. Hierarchical clustering of *Yersinia* DEGs identified a set of genes whose expression was typically high early in enterocyte pseudotime in naïve animals, but which were altered in their expression pattern by *Yersinia* infection – exhibiting either extended expression or reactivation later in enterocyte pseudotime **(Figure S6A, bracket)**. Expression of these genes was enriched in areas of higher imbalance and corresponded to a distinct *Yersinia*-driven cluster of mature enterocytes **(Figure 6C and S6B)**. We refer to these cells as “Yps enterocytes” for their origin in *Yersinia pseudotuberculosis* infection (**Figure 6D-E and S6C)** and represent a subset of all enterocytes in *Yersinia-*infected mice. To identify the genes that uniquely define this subset, we compared Yps enterocytes to all other enterocytes in *Yersinia* infection, resulting in 132 Yps-associated genes. Among these, the acute phase protein genes *Saa1* and *Saa2* were previously shown to be upregulated in epithelial cells during *Salmonella* infection^4^ and SFB colonization^63^. Other genes, including *Nos2* and *Reg3a*, encode effectors that are thought to directly limit bacterial survival and replication. Yps-defining genes were significantly associated with immune defense ontology terms, further indicating that the Yps enterocyte state might represent a direct adaptation of enterocytes to immune stimuli **(Figure 6F, Supplementary Table 4)**.

**Figure 6:**
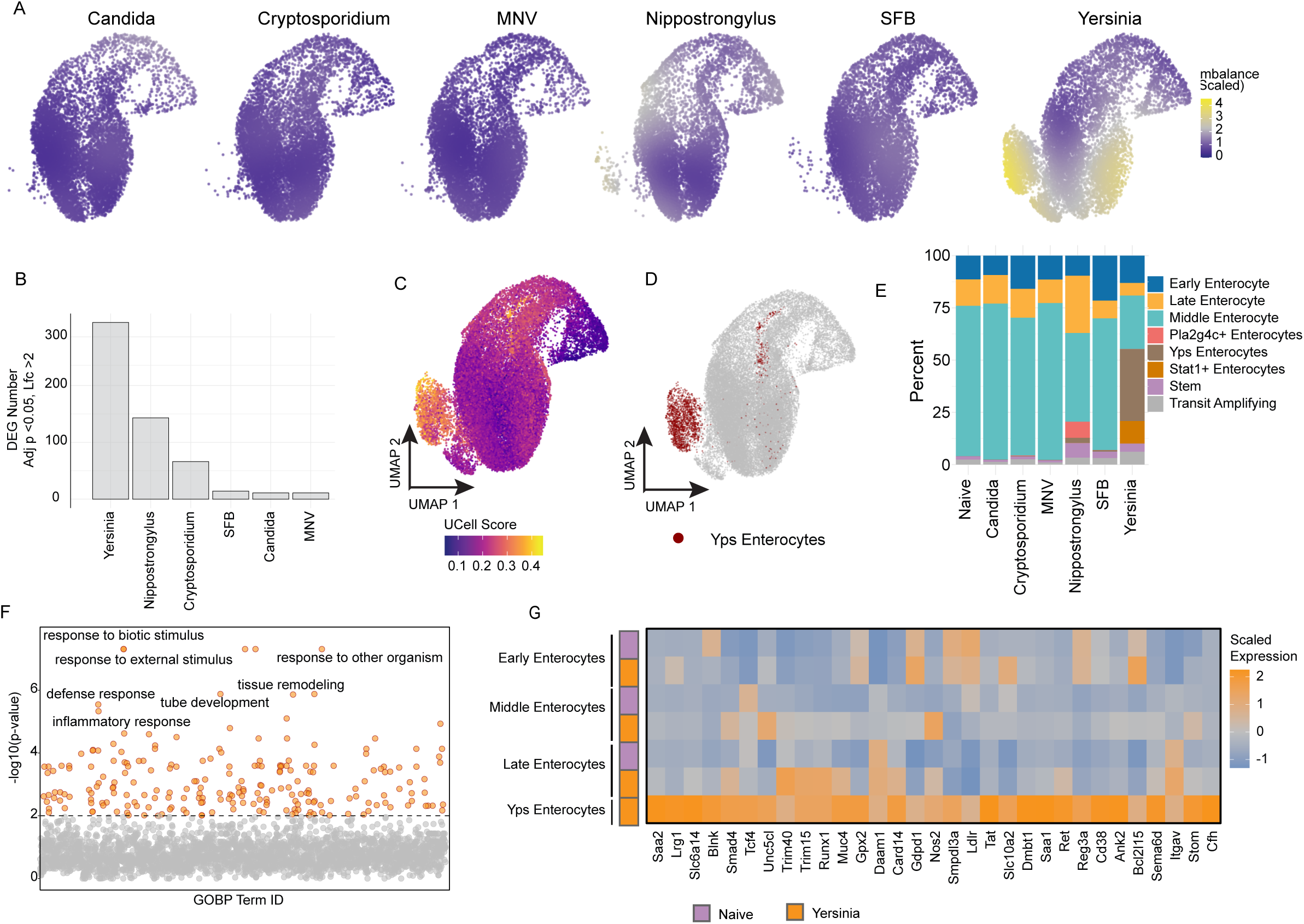
Yps enterocytes are a distinct transcriptional state. A) UMAP embeddings depicting cell imbalance scores (color) from each infection. B) DEG counts identified from pseudotime modeled comparisons of the enterocyte lineage (stem cells, transit amplifying, and enterocytes) within each infection relative to naive samples. 5000 highly variable genes included for modeling and testing, adj p < 0.05, LogFC > 2. C) UCell single cell enrichment (color hue) of select DEGs identified in enterocytes of *Yersinia*-infected samples and visualized on a UMAP dimensional reduction. D) Yps enterocytes (red) visualized on a UMAP dimensional reduction. E) Frequencies of enterocyte and stem cell subset populations (colors) within each condition. F) GO:Biological Process gene ontology results of DEGs in Yps enterocytes compared to other enterocyte populations (n = 132 genes) from *Yersinia*-infected mice. The seven most significantly enriched ontology pathways are labeled. G) Gene expression heatmap of Rcistarget-defined genes containing an enriched STAT3 motif from among Yps enterocyte-defining genes. Motifs 500bp upstream and 1000 bp downstream of the TSS were included for enrichment and enterocyte expression is shown for enterocyte populations in Naive and *Yersinia*-infected mice. Biological n: Naive = 2, Candida = 2*, Cryptosporidium = 3, MNV = 2, Nippostrongylus = 2, SFB = 2, Yersinia = 3. *Candida infection includes 1 IEC replicate.

To identify potential upstream factors and signals that might give rise to Yps enterocytes, we performed transcription factor motif enrichment using the 132 Yps enterocyte-defining genes. This analysis identified transcription factors associated with enterocyte physiology, notably *Srebf1* (NES = 4.69) (**Figure S6D)**, a regulator of fatty acid synthesis, as well as *Stat3* (NES = 3.30), a downstream effector of many cytokine-signaling cascades **(Figure 6G and Supplementary Table 5)**. The expression of genes identified as having a STAT3 motif were compared across enterocyte subsets in *Yersinia-*infected and naive mice, revealing that the STAT3 signature was specific to Yps enterocytes **(Figure 6G)**. The separation of Yps enterocytes from other mature enterocytes in *Yersinia* infected mice has interesting implications and might represent spatially restricted bacterial growth or immune signaling. Collectively, these data identify a distinct pro-inflammatory enterocyte population induced by *Yersinia* infection and potentially contributing to ongoing immune responses.

### Spatially localized inflammation during *Yersinia* infection remodels the ileal epithelium

*Yersinia pseudotuberculosis* infection results in macroscopic, dense foci of inflammatory cells in the SI called pyogranulomas (PG)^44^. PGs are highly enriched in *Yersinia* bacteria relative to the surrounding tissue and accumulate neutrophils and monocytes which utilize TNF and IL-1 cytokines to promote defense^44,64^. To investigate the relationship between Yps enterocytes and the formation of PGs, we performed spatial transcriptomic profiling of the SI from naïve and *Yersinia*-infected mice **(Figure S7A)**. We generated secondary probabilistic cell segmentations using Proseg^65^ and performed quality control filtering prior to analysis, yielding 680,410 cells across two biological replicates each from naive and *Yersinia-*infected mice **(Figure S7B-C)**. Using a panel of 5,000 gene probes we obtained an average of 566 reads per cell and 126,422 - 207,476 cells per sample **(Figure S7C)**. We then utilized GutPath as a reference to annotate this spatial data **(Figure S7D)** and observed differential clustering of cells indicative of cell state changes in the enterocyte compartment of *Yersinia*-infected mice **(Figure 7A)**.

**Figure 7:**
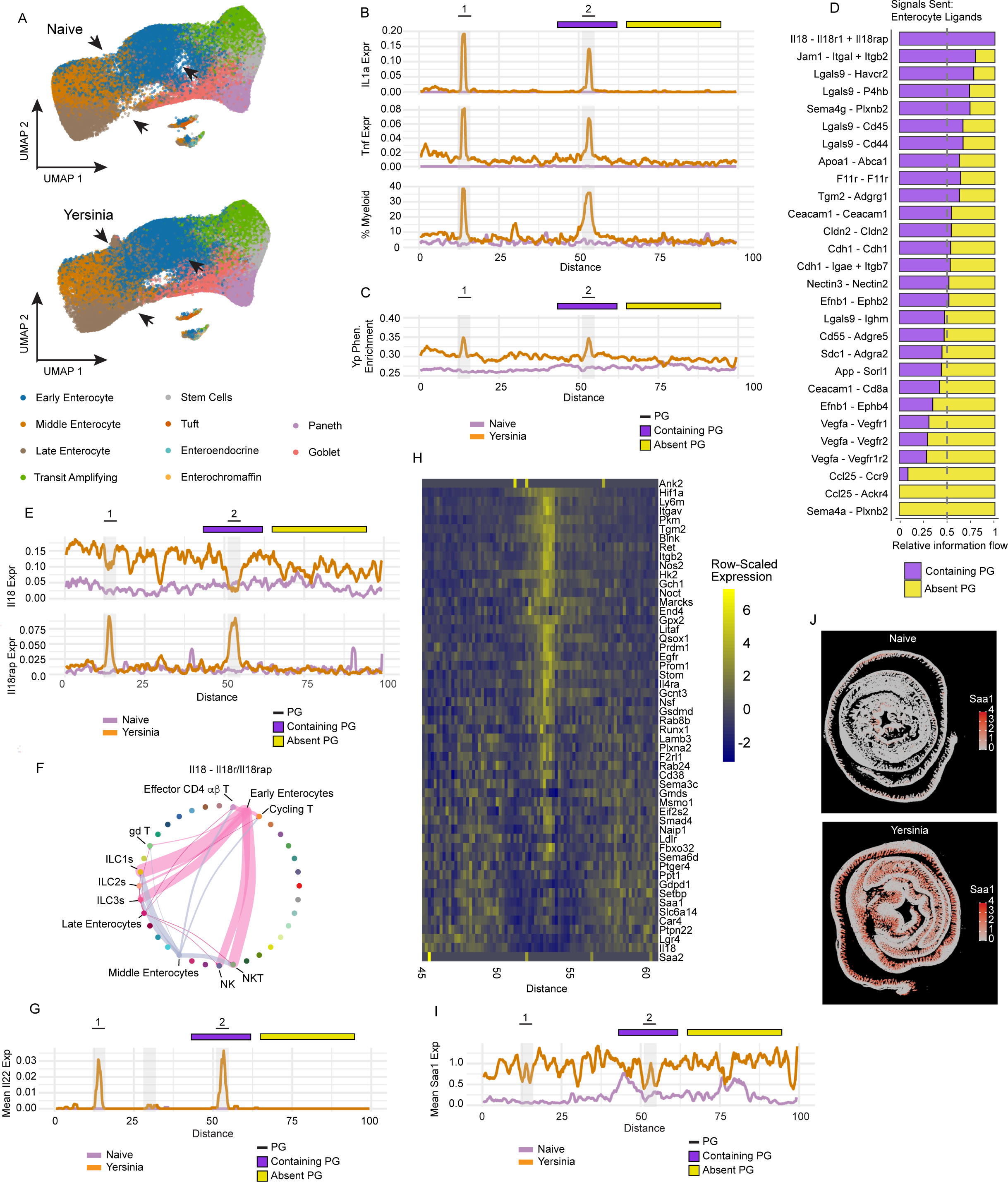
Yersinia pyogranulomas reshape the transcriptional state of surrounding enterocytes. A) UMAP dimensional reduction of subclustered epithelial cells from Naive (top) and *Yersinia* (bottom) Xenium data. Arrows indicate discrepancy between naive and *Yersinia* reductions and cell types are indicated (color). B) *Il1a* (top) and *Tnf* (middle) rolling mean expression and myeloid cell frequency (bottom) along bioinformatically informed distance (proximal to distal) in *Yersinia* and Naive spatial data. Pyogranuloma positions (shaded areas, numbered (top)) are placed with respect to “distance” within the *Yersinia* sample. Intestinal segments taken for further study of PG-containing (purple bar, top) and PG-lacking (yellow bar, top) tissue are demarcated. C) Yps enterocyte signature rolling mean UCell enrichment scores along bioinformatically informed distance (proximal to distal) in *Yersinia* and Naive samples. Pyogranuloma positions (shaded areas, numbered (top)) are placed with respect to “distance” within the *Yersinia* sample. Intestinal segments taken for further study of PG-containing (purple bar, top) and PG-lacking (yellow bar, top) tissue are demarcated. D) Cell-cell communication signaling predictions for all signals produced by enterocytes from PG containing *Yersinia* tissue (7B purple bar) relative to signals produced by enterocytes PG-lacking *Yersinia* tissue (7B yellow bar). E) *Il18* (top) and *Il18rap* (bottom) rolling mean expression along bioinformatically informed distance (proximal to distal) in *Yersinia* and Naive spatial data. Pyogranuloma positions (shaded areas, numbered (top)) are placed with respect to “distance” within the *Yersinia* sample. Intestinal segments taken for further study of PG-containing (purple bar, top) and PG-lacking (yellow bar, top) tissue are demarcated. F) Predicted cell-cell communication of IL-18 from enterocytes in PG-containing tissue to Il18 receptor components across cells. Width of interactions is proportional to the predicted strength of interactions and color indicates cell type from which IL-18 originates. G) *Il22* rolling mean expression along bioinformatically informed distance (proximal to distal) in *Yersinia* and Naive spatial data. Pyogranuloma positions (shaded areas, numbered (top)) are placed with respect to “distance” within the *Yersinia* sample. H) Gene expression of Yps-associated genes in epithelial cells along proximal to distal distance and spanning a pyogranuloma within *Yersinia* sample. Bioinformatically unrolled distance shown correspond to selected area of study (7B purple bar). I) *Saa1* rolling mean expression along bioinformatically informed distance (proximal to distal) in *Yersinia* and Naive spatial data. Pyogranuloma positions (shaded areas, numbered (top)) are placed with respect to “distance” within the *Yersinia* sample. Intestinal segments taken for further study of PG-containing (purple bar, top) and PG-lacking (yellow bar, top) tissue are demarcated. J) *Saa1* expression (color hue) overlayed on complete Naive (left) and *Yersinia* (right) tissue sections captured for spatial analysis. Xenium data are representative of 2 conditions (Naive, *Yersinia*) each with 2 biological replicates from different mice.

To understand the spatial relationships between Yps enterocytes and PGs, we identified putative PGs histologically **(Figure S7E, boxes)** and computationally ‘unrolled’ equal portions of SI Swiss rolls to create a proximal to distal placement of cells along a linearized ileum (see Methods) **(Figure S7F)**. This allowed us to reduce the spatial localization of genes to a simple linear distance across a length of SI that included two distinct PGs **(Figure 7B, top)**. Compared to naïve controls, *Il1a* and *Tnf* – two genes know to be expressed in PGs – were enriched in proximity to the PGs **(Figure 7B)**. Using the cell annotations from GutPath applied to our spatial data, we then observed peaks of myeloid cell enrichment at PG locations **(Figure 7B).** We reasoned that if the Yps enterocyte phenotype identified in our data was driven by bacterial signals, then the marker genes defining these enterocytes should localize to PGs. Cross-referencing our 132 gene Yps enterocyte signature **(Figure 6)** with the 5000 gene panel used in our spatial transcriptomics assay identified 53 genes, which we then used to test for spatially localized enrichment of the Yps enterocyte signature. We observed peaks of Yps-associated expression precisely overlapping the histologically confirmed PGs **(Figure 7C)**.

Next, to identify cell-cell signaling events that might mediate the activation of enterocytes in close proximity to PGs. We focused our analysis on a single PG and the immediately adjacent tissue **(Figure 7B, purple bar, top)** as well as a non-PG region **(Figure 7B, yellow bar, top)** and predicted cell-cell communications separately within these two distinct regions. This analysis identified ligands that were enriched in tissue closest to PGs, but not in non-PG regions of the same sample **(Figure 7D)**. Notably, this included the IL-1 family member, IL-18 and its receptors. Closer inspection of *Il18* expression showed that while it was excluded from the region immediate atop the PG, it was highly expressed in the adjacent tissue **(Figure 7E, top),** and the *Il18rap* receptor was highly expressed directly overlapping PGs, suggesting that enterocyte IL-18 production may signal to inflammatory cells residing in the PG. Consistent with this notion, our cell-cell communication analysis predicted early enterocytes to be the main producers of IL-18, which can signal to NK and NKT cells, as well as ILC1 and ILC3 cells **(Figure 7F)**. We previously demonstrated a STAT3-associated gene expression pattern in Yps enterocytes and here have shown the regulation of *Il18* proximal to PGs. We asked whether IL-22 – a STAT3-activating cytokine with pleotropic effects, including inducing enterocytes to produce IL-18^66^ – was increased at PGs. Indeed, we found that *Il22* expression peaked at the site of PGs in the *Yersinia-*infected mice **(Figure 7G)**.

Although Yps-associated expression peaked within PGs, it remained elevated above naïve control well beyond the location of PGs **(Figure 7C)**, suggesting that individual enterocyte genes may vary spatially in their expression during infection. To examine this in more detail we focused on the region spanning the second PG and including the immediately adjacent tissue **(Figure 7B, purple bar, top)** and viewed expression of each of the 53 enterocyte genes as a heatmap across that region **(Figure 7H)**. This analysis showed that most of these genes – including *Hif1a*, *Nos2*, and the integrins *Itgav* and *Itgb2* – showed highly restricted expression positioned immediately atop the PG in the enterocyte compartment **(Figure 7H)**. However, *Saa1* remained elevated across the entire region we examined, compared to naïve control **(Figure 7I)**, and was induced in enterocytes across the entire ileum during infection **(Figure 7J)**. These data identify a *Yersinia pseudotuberculosis-*associated enterocyte phenotype that is spatially linked to the formation of pyogranulomas in the small intestine, but which has reverberations in the gene expression of enterocytes extending far beyond sites of infection.

## Discussion

Infections of the small intestine are major causes of malnutrition, growth stunting, and death globally. Leveraging intestinal pathogens/commensals and experimental infections we created GutPath (available at GutPath.org), a resource that comprises RNA and protein data for over 300,000 single cells (91 cell states) from the ileum and nearly 200,000 cells (49 cell states) from the ileal-draining MLN, across six different infections and naive controls. Responses to fungal, bacterial, parasitic helminth and protozoan, and viral model organisms are represented and capture diverse immune states spanning Th1, Th2, and Th17-dominant immune archetypes, with all major cell populations and many rare cell types profiled. The breadth and depth of GutPath allows investigators to ask new questions about conserved patterns of biology in the ileum, to contextualize new and existing mucosal datasets, and to better interpret mucosal immune responses in other infectious or inflammatory diseases.

A major impediment to the efficient analysis of new single cell datasets is accurate and detailed cell annotation, particularly in extremely dynamic and complex tissues such as the gut. Although multiple resources exist to aid in this process^34,67^, they do not reflect diverse immune cell activation states, particularly those initiated by infection, thus requiring investigators to undertake arduous manual annotation efforts. GutPath addresses this problem by providing multiple layers of cell state annotations, from broad cell lineages to highly specific cell phenotypes, each with accompanying marker profiles. In addition to all raw sequencing data and code being publicly accessible, gene expression in different cell types and within different pathogen model systems contained in GutPath can be directly queried in a web browser via the CELLxGENE^68,69^ Visualization In Plugin (VIP)^70^, available at GutPath.org. This atlas also provides a critical foundation to leverage rapidly evolving large language models to explore complex cellular data. Such approaches streamline analysis workflows, significantly improving research efficiency while accelerating the translation of high-resolution datasets into potentially actionable insights.

This atlas is a rich resource for understanding enterocyte biology, particularly because our sample preparation yielded the complete developmental ‘lifespan’ of an enterocyte, ranging from crypt-resident stem cells to transit amplifying cells, to mature absorptive enterocytes present along the length and tip of the villus **(Figure 2)**. This allowed us to use pseudotime-based trajectory analyses to assign transcripts to precise enterocyte stages of development, revealing a highly structured pattern of expression for 51 nutrient and fluid transporters from the crypts to the tips of the villi, with 20 of these genes showing significant dysregulation during one or more infection **(Figure 4C-D)**. Disruption of the intestinal epithelium in Crohn’s disease patients and during infection can lead to a wide range of malabsorption phenotypes, but the molecular mechanisms underlying this clinical manifestation are poorly understood. The infection-dependent dysregulation of key transporters observed in this study constitutes potential molecular mechanisms for pathogen disruption of ileal biology, and merits further investigation. In addition to enterocytes, our atlas captured immune cells interspersed within the epithelial layer, as well as in the underlying lamina propria and draining lymph node, thus empowering researchers to predict cell-cell communication occurring between immune cells or between immune and stromal cells. This is illustrated by our identification of a unique enterocyte phenotype induced by *Yersinia* infection **(Figure 6)** which our data suggests may be driven by IL-22/STAT3 signaling to produce IL-18, that then signals through IL18rap on ILCs, NK cells, and NKT cells residing in the *Yersinia* pyogranuloma.

In addition to examining transcriptional responses along enterocyte pseudotime, we also explored how mucosal immune responses were influenced by proximity to tissue pathology and pathogen load **(Figure 7)**. Our experiments with *Yersinia pseudotuberculosis* revealed that some enterocyte transcriptional states co-localized precisely with tissue pathology (pyogranulomas), while other states were more regional and found adjacent to pyogranulomas, and still other states were evident across the whole tissue. Similarly, our experiments with *Nippostrongylus brasiliensis* which primarily infects the proximal small intestine, showed dramatic metabolic reprogramming of the epithelium in the distal small intestine **(Figure 5)**. Taken together, these results reveal a ‘ripple effect’ of gene expression that can emanate from areas of the tissue where pathogen load and pathology are highest, thus suggesting that even highly focal or well-contained infections may still influence enterocyte behavior far from the site of tissue insult. Since our experiments only covered a single, early (< 1 week) timepoint for each infection, it is unclear the extent to which these transcriptional programs are sustained and how they impact the development of effector or memory T cell responses. Nevertheless, this finding has important implications not just for GI infections, but also for inflammatory diseases like Crohn’s disease where lesions form in the small intestine and are a central part of disease pathogenesis. Our spatial data provide a granular view of the genes most closely linked to tissue pathology and their relative localization or spread across the tissue, which should be investigated as therapeutic targets and drivers of tissue remodeling and bacterial control.

Even though immune cell infiltrates and proinflammatory gene expression were concentrated around pyogranulomas in *Yersinia*-infected mice, we showed that *Saa1* was widely expressed in enterocytes across the entire distal SI. SAA1 is known to be induced by IL-1 family members and TNF- , has direct antimicrobial properties^71,72^, and promotes intestinal epithelial cell migration during wound healing^73^. However, SAA1 can also contribute to proinflammatory responses through the generation of pathogenic Th17 cells^74^ and has been linked to sites with active inflammation in Crohn’s disease patients^75^. It is possible the highly focal nature of proinflammatory cytokine induction around pyogranulomas is sufficient to elicit tissue-wide expression of downstream targets. Conversely, there may be a mechanism by which enterocytes propagate inflammatory signals across large distances from an initial inflammatory insult. Delineating how inflammatory signaling events unfold over time and space to modulate gene expression in enterocytes can have important repercussions for identifying new therapeutic opportunities for GI diseases.

## Methods

### Mouse care and use

All animal procedures and mouse husbandry were performed under guidelines and approval from the University of Pennsylvania Institutional Animal Use and Care Committee (IACUC protocol numbers 805432, 804523, and 807356). Female specific pathogen free C57BL/6 mice aged 4-7 weeks were purchased from Jackson Laboratories and housed at the University of Pennsylvania Animal Care Facilities 1-2 weeks prior to study. All mice were free of Segmented Filamentous Bacteria (SFB) colonization prior to experiments.

### Infections

#### Cryptosporidium parvum

Mouse adapted *Cryptosporidium parvum* expressing nanoluciferase was propagated as described previously in *Ifng*^−/−^ mice^76^. Oocysts were purified from the feces of infected mice by use of sucrose flotation and cesium chloride gradients. Experimental mice aged 6-7 weeks were infected with 5×10^4 oocytes by oral gavage. Infection was confirmed and fecal shedding of oocysts was measured by luminescence (Glomax reader, Promega). Mice were euthanized at 4 days post-infection (dpi) for tissue collection.

#### Nippostrongylus brasiliensis

Third stage (L3) *Nippostrongylus brasiliensis* were purified and washed in PBS. Mice were injected subcutaneously with 200 infective L3 in PBS and euthanized 5 dpi and tissues collected. A second cohort of mice were simultaneously administered 200 L3 and burden was measured by worm count within the small intestine at 6 dpi to ensure L3 viability and infectivity.

#### Yersinia pseudotuberculosis

*Yersinia pseudotuberculosis* (clinical isolate strain 32777, serogroup O1) was expanded at 28°C for 16 hours in 2xYT broth supplemented with 2 ug/ml triclosan (Millipore Sigma). Mice were fasted for 16 hours and administered 2×10^8^ colony-forming units (CFU) of *Yersinia* in 200 μL PBS as previously described^44,64^. Mice were euthanized and tissues collected at 5 dpi. Spleens from infected mice were collected in sterile PBS, weighed, and homogenized for 40 seconds with 6.35 mm ceramic beads (MP Biomedical) using a FastPrep-24 bead beater (MP Biomedical). Tissue homogenates were serially diluted 10-fold in PBS, plated in triplicate on LB agar supplemented with 2 ug/mL triclosan, and incubated for 2 days at room temperature. Countable colonies at the lowest dilution step were used to calculate the CFU per gram.

#### Murine Norovirus

Molecular clones of the MNV strain CR6 were generated as previously described to produce working stocks of infectious virus^77^. Plasmids encoding the viral genome were transfected into 293T cells (ATCC) to generate a P0 stock. This P0 stock was subsequently amplified through two passages in BV2 cells at a multiplicity of infection (MOI) of 0.05 to produce P2 stock. For viral stock preparation, infected BV2 cells were freeze-thawed, cell lysates and supernatants were centrifuged at 1,200 × g for 5 minutes. Supernatants were then filtered through a 0.22-μm filter and concentrated using 100 kDa MWCO Amicon Ultra centrifugal filters. Viral titers were determined by plaque assay using BV2 cells overlaid with DMEM (Corning) containing 1% methylcellulose (Sigma-Aldrich). Plaques were evaluated three days post-infection by crystal violet staining. Mice aged 7 weeks were infected orally with 3×10^6^ PFU and euthanized at 7 dpi for tissue collection.

#### Candida albicans

A single colony of *Candida albicans* CHN1 was cultured in 5 ml of Sabouraud dextrose media at 30°C for 16 hours. Cells were pelleted by centrifugation at 1,000 × g for 5 minutes and resuspended in 2 mL of PBS. At 6-7 weeks of age, mice were infected by oral gavage of 5×10^7^ CFU and were euthanized at 7 dpi for tissue collection.

#### Segmented Filamentous Bacterium (SFB)

Fecal pellets of segmented filamentous bacteria (SFB) colonized mice were collected and stored at - 80°C. Primary SFB colonization was established in 6-7 week-old SFB-negative mice by oral gavage of homogenized SFB-containing fecal pellets as previously described^78^. Mice were euthanized and tissues collected at 7 dpi for analysis. Successful colonization was measured at day 5 post gavage by qPCR targeting of SFB.

### Tissue processing

All samples were handled on ice and centrifugation performed in pre-chilled 4°C centrifuges to promote cell viability. The ileal-draining mesenteric lymph node was removed and placed in PBS prior to dissociation. Mechanical and chemical dissociation were utilized including 20 minutes incubation at 37°C in 700 uL digest buffer (Complete RPMI (RPMI+10% FBS + 0.1% β-mercaptoethanol + 1% non-essential amino acids + 1% sodium pyruvate + 1% penn/strep) + 0.5mg/mL DNase + 0.167 mg/mL Liberase TL). The distal third of the small intestine was isolated, Peyer’s patches were excised and discarded, and the tissue was minced. The epithelial layer was separated following incubation/shaking of intestinal tissue at 37°C for 20 minutes in 20 mL of chelating and reducing buffer (1L HBSS + 5% FBS + 5mM EDTA ) + 0.154 mg/mL DTT. Further mechanical shaking and washes were performed to separate and collect the epithelial layer. The lamina propria was then digested in 20mL of digest buffer for 30 minutes at 37°C, shaking. All samples were filtered through 70 um and 40 um filters. Dead cell removal was performed for all IEC suspensions and for LP samples where indicated by viability (Miltenyi Biotec #130-090-101)

### Single cell sequencing

Single cell suspensions were processed according to 10x Genomics and Biolegend protocols to create Cellular Indexing of Transcriptomes and Epitomes (CITE-seq) libraries. In brief, cells were stained with the TotalSeq B Mouse Universal Cocktail V1.0, which includes oligo-conjugated antibodies for ∼100 surface proteins (Biolegend). Cells and 10x Genomics barcoded gel beads were then encapsulated using GEM-X 3’ chips and a Chromium Controller device. Standard library amplification and indexing were performed with either 10x Genomics GEM-X single cell 3’ v3 or v4 chemistry. All paired-end libraries were sequenced on a NextSeq 2000.

### Analysis of single cell data

#### Data preprocessing and integration

Cell Ranger (v8.0.0) was utilized with mouse genome GRCm39 to generate raw and filtered feature-barcode matrices. Matrices were imported into R (v4.5.0) using Seurat (5.3.0) and underwent primary filtering to retain cells that met the following criteria: RNA and Antibody derived tag (ADT) library sizes within ±2.5 median absolute deviations of the median, >750 detected genes, and <5% mitochondrial RNA content for the MLN and <20% mitochondrial content for Ileum samples. ADT counts were normalized using the DSB (denoised and scaled by background) method (dsb v2.0.0)^79^ with isotype control features included for technical correction. Merged and processed Seurat objects were used for all downstream analyses.

RNA and ADT data were independently processed, integrated, and visualized. Standard variable feature identification (n = 3000), normalization, and scaling were applied. Independent samples were integrated via Harmony using both Louvain and Leiden clustering (Monocle3 v1.3.1). RNA and ADT modalities were incorporated through WNN analysis for further clustering and subsequent dimensional reduction. Doublet removal was performed using scDblFinder (v1.22.0) and regression of cell cycle expression^80^ was performed prior to final dimensional reduction visualization.

Clusters and cells were initially annotated using both automated reference-based methods (SingleR v2.11.1, reference = ImmGenData) and manual expert annotation through applied literature search. Subsequent annotation relied on label transfer to newly added data and manual refinement. For most major populations, sub-setting, re-clustering, and the generation of new dimensional reductions was performed to describe and visualize cellular heterogeneity and perform detailed annotation as seen in Figure 2 and Figure S3.

#### DEG analysis using MAST

Differential gene expression between infection conditions and the naive states was calculated separately for each cell type at an intermediate resolution level, restricted to comparisons with >10 cells per condition. The Seurat (5.3.0) deployment of Model-based Analysis of Single-cell Transcriptomics (MAST) is robust to drop out and was used with the following latent parameters: genes per cell, UMI per cell, percent ribosomal reads, and percent mitochondrial reads^54^. To control the false discovery rate (FDR), p values were globally adjusted (Benjamini–Hochberg) and a permissive FDR of <0.05 was utilized for data exploration.

#### Gene ontology and Enrichment analysis

Select gene collections identified in differential expression analyses and cross-infection comparisons were utilized for gene ontology (gProfiler v0.2.3). The gene ontology biological process (GOBP) catalogue was used in most instances with FDR correction. Ontologies containing fewer than 20 or more than 500 genes were excluded. To calculate scoring and enrichment of gene sets at the single cell level we utilized the UCell method as it is scalable and robust to gene drop out (escape v2.4.0 & UCell v2.6.2). Further scaling of gene set scores was applied for visualization where indicated.

#### Metabolic Analysis

To infer metabolic activity states across single cells, we applied scCellfie and the included metabolic task database (v0.4.7)^59^ (Python v3.10.18, scanpy v 1.11.3). Individual task scores were calculated with the following parameters: smooth cells = True, alpha = 0.33, n nearest neighbors = 10, and batch indicator = Mouse. Normalized cell type-specific metabolic activity was summarized by computing the trimean of task scores within each cell type and visualized in radial plots across infection conditions. Differential analysis of metabolic activity was calculated for each cell type relative to the naive samples and significance was determined at FDR <0.05, Cohen’s D > 0.5 and Log2FC >0.585 (1.5 FC). Significantly altered lipid metabolic pathways identified in enterocytes from *Nippostrongylus*-infected mice were visualized.

#### Pseudotime and trajectory-based differential expression

To model epithelial differentiation along the crypt–villus axis, ISCs, transit amplifying cells, and enterocytes were subset from the data and re-clustered with new dimensional reductions. Pseudotime trajectories were inferred using Monocle3 and ISCs were manually selected as the root node^81^. Within the naive condition, pseudotime-associated genes were identified using a generalized additive model (GAM) and trajectory-based differential expression analysis tradeSeq (v1.13.12) to test for significant expression changes as a function of pseudotime^55^. Genes with adjusted p < 0.05 (Benjamini–Hochberg correction) were considered significantly dynamic and filtered for pre-defined genes associated with solute transport.

To measure infection-induced remodeling of epithelial differentiation, the same pseudotime ordering was used to compare expression patterns between naive and infection conditions. Condition-specific pseudotime expression of previously identified pseudotime-associated metabolic and solute transport genes were selected for visualization. General trends in infection-directed pseudotime expression were analyzed using the pseudotime scores as a continuous covariate in a linear model framework, following the same approach. Genes exhibiting significant condition-by-pseudotime interactions were classified as differentially regulated along the trajectory. Differentially expressed genes in this model from enterocytes of Yersinia infected mice were further studied by gene ontology and functional analysis.

#### Single-cell analysis: transcription factor activity inference

To identify transcription factors regulating the Yps enterocyte transcriptional module, motif enrichment was performed using the RcisTarget package (v1.28.1)^82^. The Mus musculus motif ranking database (mm10 refseq-r80 500bp_up_and_100bp_down_tss.mc9nr.genes_vs_motifs.rankings.feather) and the corresponding motif-to-transcription factor annotation table (motifs-v9-nr.mgi-m0.001-o0.0.tbl) were downloaded from the cisTarget resource repository. Motifs with normalized enrichment scores (NES) > 3.0 were considered significantly enriched and were annotated to candidate TFs. Motif-gene relationships of selected enriched motifs were then visualized.

#### Cell-cell communication analysis

Cell-cell communication analysis was performed using CellChat (v2.1.2)^83^. Separate CellChat objects were created from Seurat-processed ileal infection datasets. Ligand–receptor interactions were inferred from receptor ligand pairs contained in the CellChatDB.mouse database. Identification of overexpressed genes and interactions, computation of communication probabilities (bootstrap n = 50), and aggregation at the signaling pathway level were also performed. Networks were compared across conditions using merged conditions and signaling centrality scores were calculated. Visualization focused on RankNet plots (significance threshold p < 0.05) to identify condition-specific signaling changes in late enterocytes and on IL18-related signaling networks to highlight *Yersinia*-associated communication alterations.

### Spatial transcriptomics

Spatially resolved transcriptomic profiling was carried out using the 10x Genomics Xenium platform and the pre-designed Xenium Prime 5K Mouse Gene Expression Panel. Infected and naive mice were euthanized and the small intestine isolated and flushed before 5 cm of the terminal ileum were isolated and Swiss rolled. Tissues were fixed in 10% neutral buffered formalin for 18-24 hours prior to paraffin embedding. Sectioning and Xenium imaging were performed in partnership with the University of Pennsylvania Dental School Spatial Transcriptomics Core.

Cell segmentation of Xenium data was performed using Proseg (v2.0.3), a probabilistic cell segmentation framework^65^, and Xenium Ranger (v3.11). For each sample, raw Xenium outputs generated by Xenium Ranger were imported and cell-level quality metrics, including total transcript counts and detected features, were used for filtering. Cells with values outside ±2.5 median absolute deviations (MAD) from the median for either metric were excluded. The remaining high-quality cells were retained for downstream SCT normalization, dimensional reduction, and clustering.

To linearize the spatial organization of the intestinal Swiss roll, Xenium spatial coordinates were exported and processed using IntestLine to manually define points along the outer and inner boundaries of the rolled tissue^84^. These boundary coordinates were used to construct a closed polygon using the sf (1.0-21) package in R. Cells located within the polygon were subset from the data and a continuous spatial trajectory (centerline) through the tissue was calculated. Cumulative Euclidean distances were then computed along this line to establish a normalized positional coordinate representing the cell’s relative proximal to distal location along the unrolled axis. Each cell within the polygon was assigned a corresponding value based on the nearest point on the interpolated centerline. This derived spatial variable was added to the Seurat object and used for downstream visualization and analysis, effectively “unrolling” the Swiss roll into a one-dimensional axis.

## Data availability

CITE-seq data are available as processed, fully annotated Seurat objects, and as downsampled data loaded into web-accessible CellxGene instances at GutPath.org. In addition, all raw fastq files for CITE-seq experiments are available on the Sequence Read Archive (SRA accession, PRJNA1378118). Xenium 5K mouse spatial transcriptomic data are available on the Gene Expression Omnibus (GEO accession, GSE313636).

## Code availability

All code used in analysis and visualization of CITE-seq and spatial transcriptomic data is available on Github at: https://github.com/hartandrew/GutPath

## Contributions

Planning and project management: A.H., M.M., C.H., B.E.H., I.S.C., Y.Y., K.C., J.M., H.Y., X.H., D.C., I.I.I., I.E.B., D.A., C.A.H., D.P.B. Infections: M.M., C.H., B.E.H., I.S.C., S.B., Y.Y., H.Y., X.H. Tissue/single cell suspension collection: A.H., M.M., C.H., B.E.H., I.S.C. 10X CITE-seq sequencing: D.C. Tissue collection for spatially resolved transcriptomics: S.B., B.W. Single cell and spatial transcriptomic analysis: A.H., R.X. Computational resource management: R.X., A.H., D.P.B. Study conceptualization: The MIST Consortium, B.S., D.A., C.A.H., D.P.B. Intellectual contribution: A.H., K.C., J.M., I.I.I., B.S., S.S., I.E.B., D.A., C.A.H., D.P.B. Writing of the manuscript: A.H., C.A.H., D.P.B.

## Funding

This research was supported by the National Institutes of Health (DK126871, AI151599, AI095466, AI095608, AR070116, AI172027, DK132244 all to D.A., K99AI180354 to H.Y.), the Allen Discovery Center program, a Paul G. Allen Frontiers Group advised program of the Allen Family Philanthropies, the Crohn’s and Colitis Foundation Research Fellowship Award (#937437 to H.Y. and #1455492 to X.H.), Cure for IBD, Weill Cornell Medicine Jill Roberts Institute and the Sanders Family, the Rosanne H. Silbermann Foundation and the Parker Institute for Cancer Immunotherapy at Weill Cornell Medicine (all to D.A.).

## Supplementary Tables

**Supplementary Table 1:** Marker genes that define coarse and intermediate cell state annotations for MLN and ileum. Separate tabs are shown for ‘fine’ (most refined) cell state annotations separated by Epithelial, B cell, T cell, stromal, and myeloid cell populations.

**Supplementary Table 2**: Marker genes from Supplementary Table 1 further condensed using Random forest-based machine learning approach to generate necessary and sufficient marker profiles for each cell state and level of annotation. One tab each for coarse, intermediate, and fine level annotations.

**Supplementary Table 3**: 134 genes known to be involved in regulation of metabolism and nutrient transport in the intestine. This list was cross-referenced with our 4,278 enterocyte DEGs to identify the 51 genes shown in Figure 4G and the 20 genes shown in Figure 4H.

**Supplementary Table 4**: Gene Ontology (GO) enrichment (Biological Process) results for 132 Yps-associated genes identified by comparing Yps enterocytes to all other enterocytes in *Yersinia* infection. Results are depicted in Figure 6F

**Supplementary Table 5**: Transcription factor motif enrichment results generated using the 132 Yps enterocyte-defining genes. Results depicted in Figure 6G and Figure S6D.

## Notes

https://gutpath.org/

### Competing Interest Statement

The authors have declared no competing interest.

