## Supplementary Figure 1 for "Diverse infections transcriptionally reprogram the intestinal epithelium and epithelial-immune cell interactions"

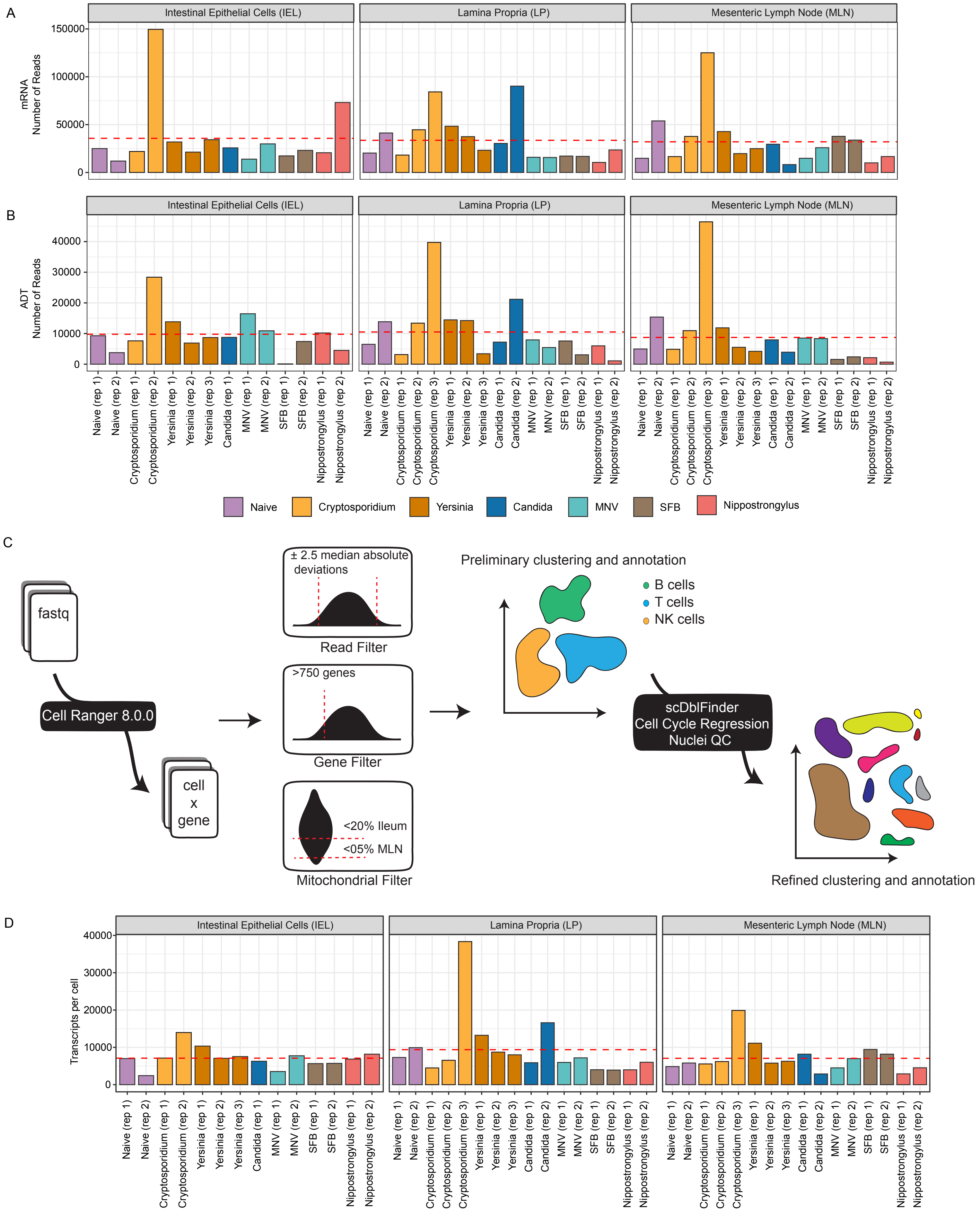


Figure S1: GutPath quality control and processing. A) Raw sequenced mRNA reads (mean) for each mouse across the separate libraries: IEC, LP, and MLN. B) Raw sequenced ADT reads (mean) for each mouse across the separate libraries: IEC, LP, and MLN. Red dotted lines indicate averages across samples. C) Bioinformatic workflow schematic showing data filtering cutoffs and major quality control measures. D) Filtered mRNA UMIs (mean) for each mouse across the separate libraries: IEC, LP, and MLN. Colors indicate infection.
