## Supplementary Figure 2 for "Diverse infections transcriptionally reprogram the intestinal epithelium and epithelial-immune cell interactions"

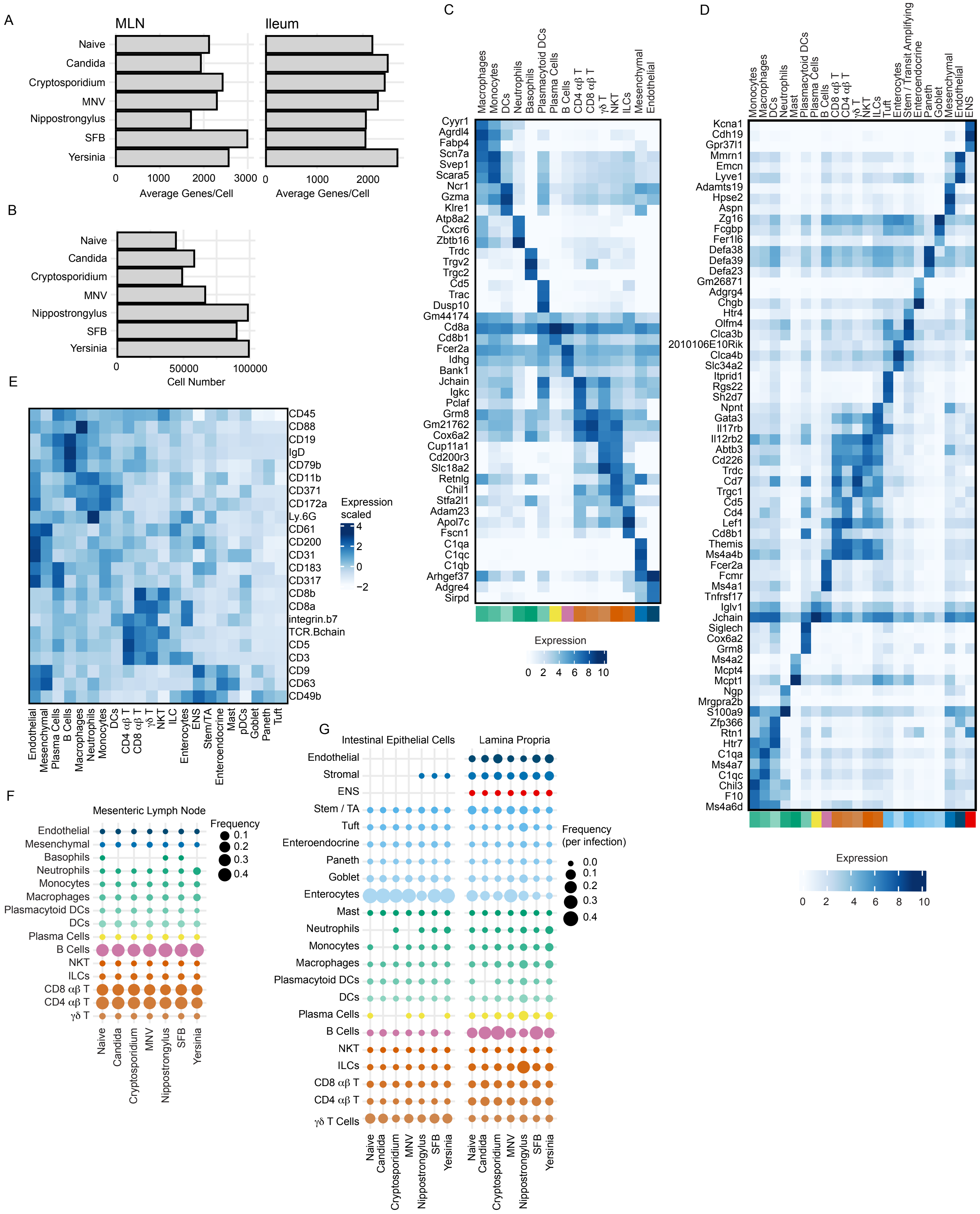


Figure S2: GutPath annotation. A) CITE-seq mean genes per cell captured across all conditions in the MLN (left) and Ileum (right). B) Total numbers of high-quality filtered cells obtained across all replicates for each infection model. C) Normalized gene expression (color hue) for select major cell types (annotation bar color) in the MLN, top 3 genes per cell type are shown. D) Normalized gene expression (color hue) for select major cell types in the ileum, top 3 genes per cell type are shown. E) Scaled expression (color hue) of cell type marker proteins in the ileum, highly differentiating proteins selected. F) Frequencies of selected major cell types for each condition in the MLN. G) Frequencies of selected major cell types for each condition in the IEC (left) and LP (right). Biological n: Naive = 2, Candida = 2*, Cryptosporidium = 3, MNV = 2, Nippostrongylus = 2**, SFB = 2, Yersinia = 3. *Candida infection includes 1 IEC replicate. **Nippostrongylus biological replicates include both ileal and duodenal-derived LP and IEC samples.
