## Supplementary Figure 3 for "Diverse infections transcriptionally reprogram the intestinal epithelium and epithelial-immune cell interactions"

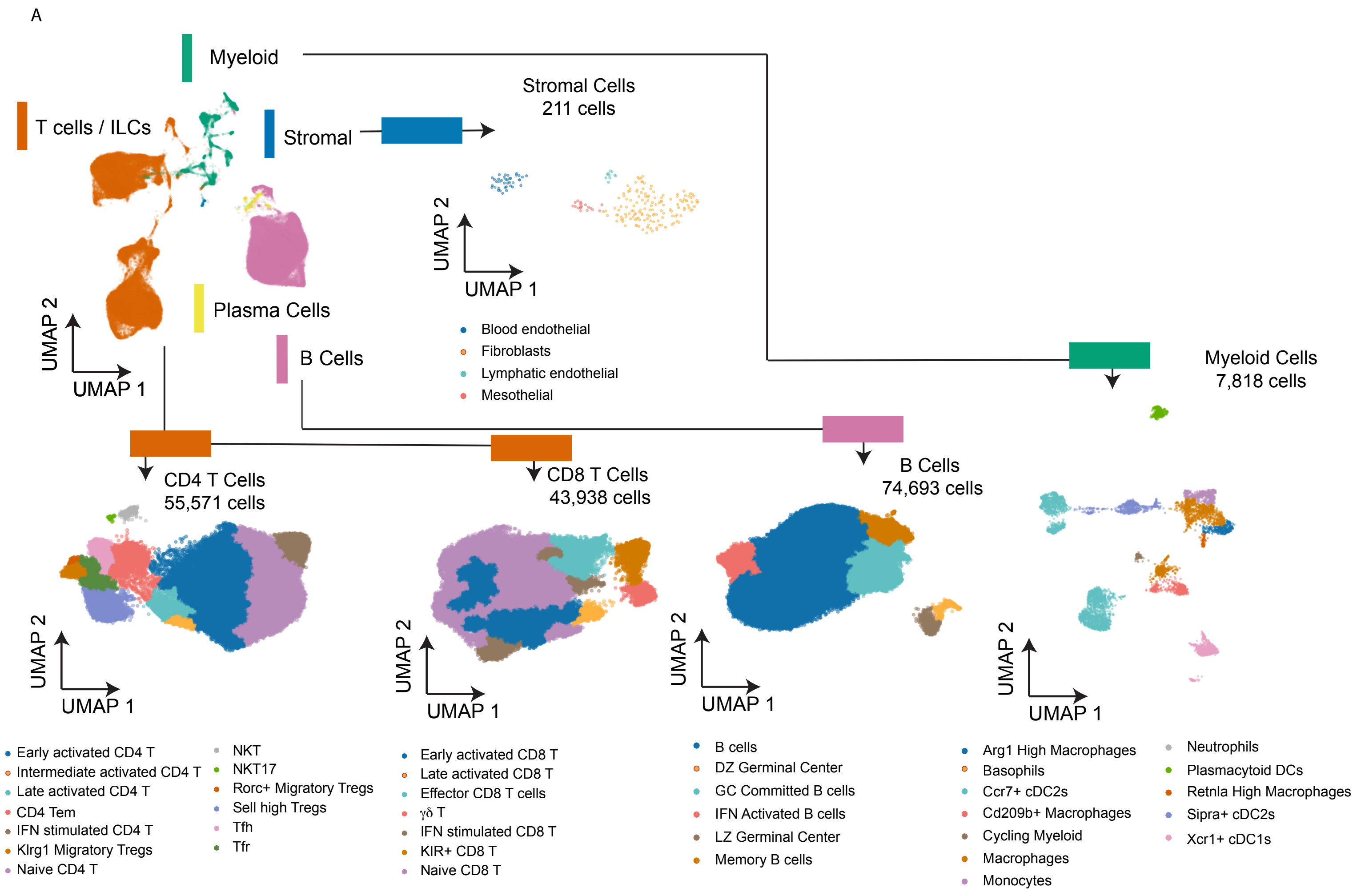


Figure S3: GutPath refined annotation. A) Schematic UMAP diagrams showing total MLN data and major subsets that were segregated, re-clustered, and subjected to refined cell type annotation. Cell numbers and refined cell state annotations of each subset are provided. Major cell lineages to which each subset belongs are indicated (bar) by color. Plasma cells were not further subset and no subset clustering is shown. Biological n: Naive = 2, Candida = 2*, Cryptosporidium = 3, MNV = 2, Nippostrongylus = 2**, SFB = 2, Yersinia = 3.
