## Supplementary Figure 4 for "Diverse infections transcriptionally reprogram the intestinal epithelium and epithelial-immune cell interactions"

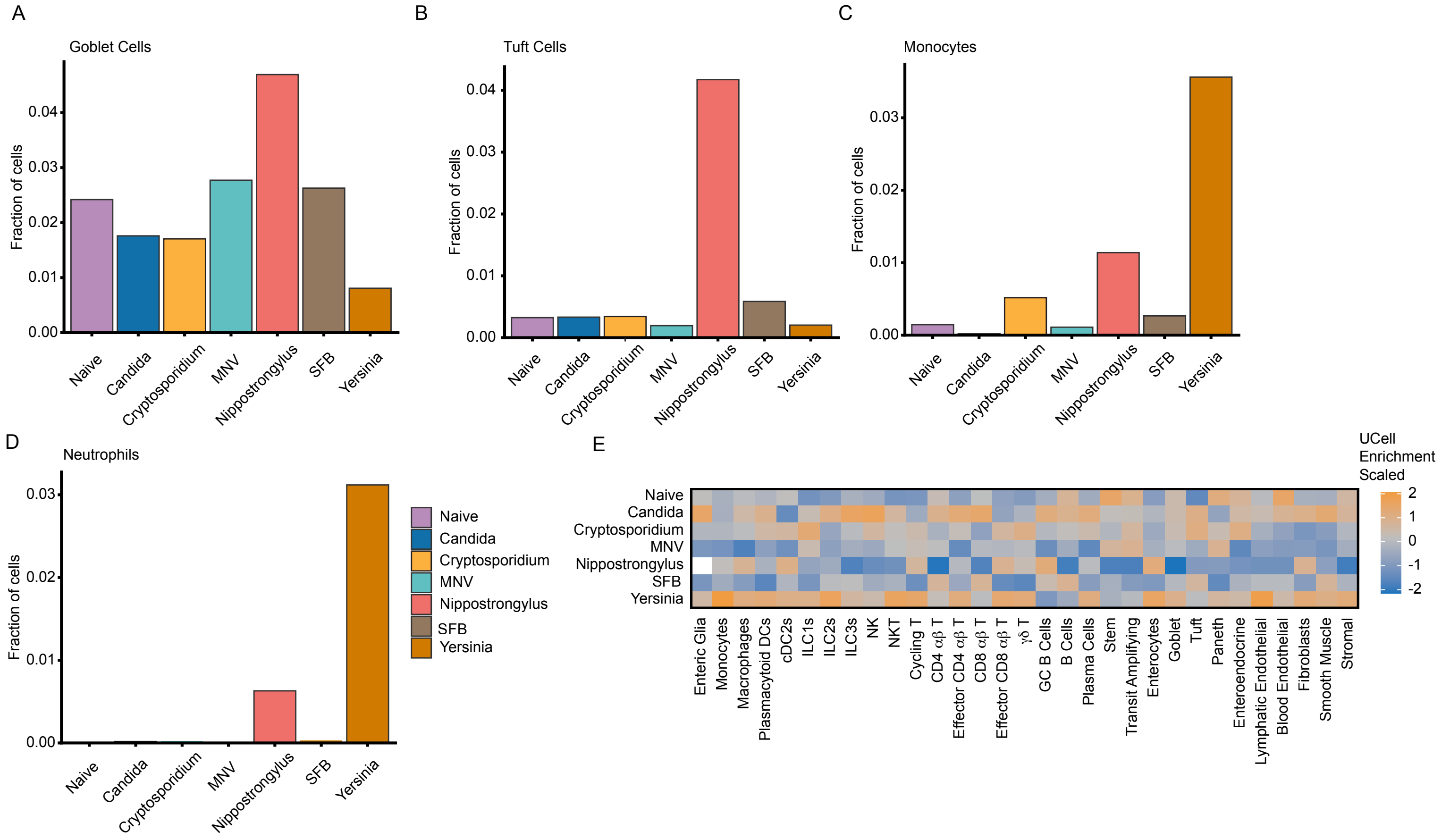


Figure S4: GutPath captures predicted immune architecture. A) Frequencies of goblet cells in the ileum for each infection (color). B) Frequencies of tuft cells in the ileum for each infection (color). C) Frequencies of monocytes cells in the ileum for each infection (color). D) Frequencies of neutrophils in the ileum for each infection (color). E) Scaled gene set enrichment score (UCell method) per cell type and infection of the Hallmark Hypoxia gene set. Biological n: Naive = 2, Candida = 2*, Cryptosporidium = 3, MNV = 2, Nippostrongylus = 2, SFB = 2, Yersinia = 3. *Candida infection includes 1 IEC replicate.
