## Supplementary Figure 5 for "Diverse infections transcriptionally reprogram the intestinal epithelium and epithelial-immune cell interactions"

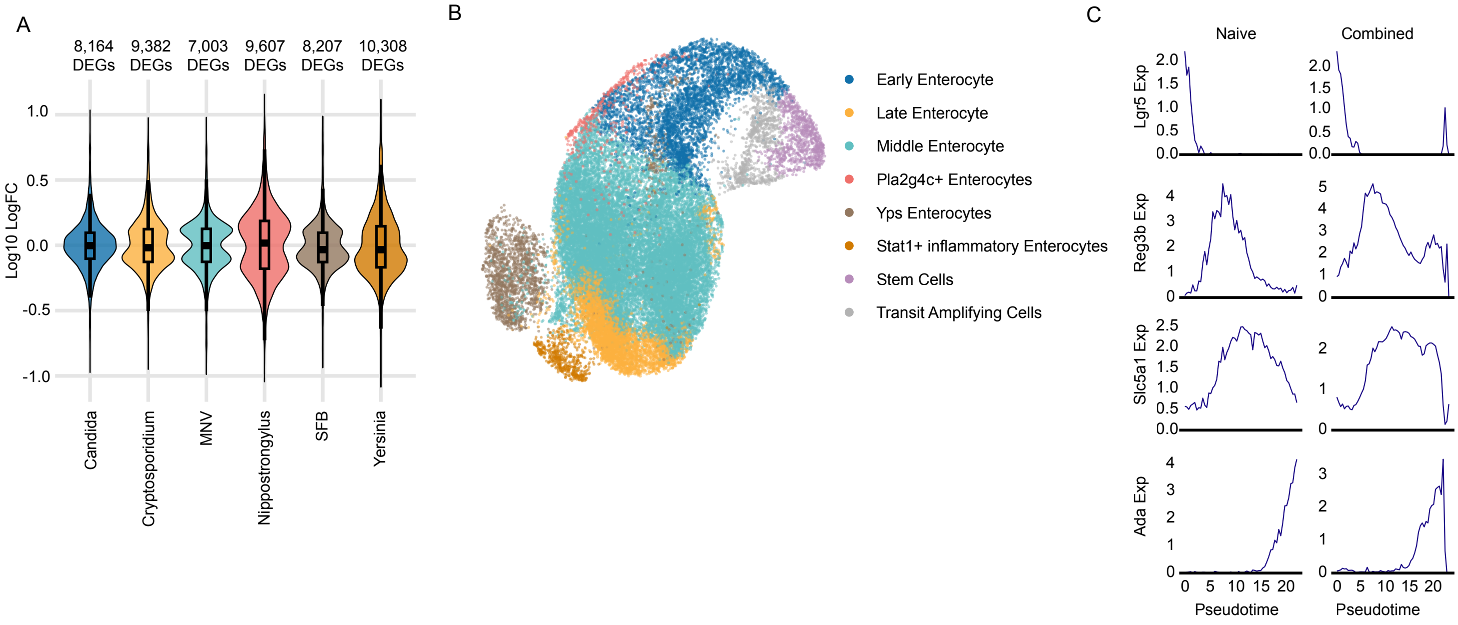


Figure S5: GutPath demonstrates infection-driven enterocyte changes to transcription. A) MAST differential expression analysis results for each infection (color), adjusted p <0.05. Log scaled logFC is shown. B) UMAP visualization constructued from stem cells, transit amplifying cells, and enterocytes across all infections. Cell state annotations are marked by color. C) Expression of *Lgr5*, *Reg3b*, *Slc5a1*, and *Ada* across pseudotime in the enterocyte and stem cell compartment of naive mice (left) or the combined infection/colonization data sets (right). Biological n: Naive = 2, Candida = 2*, Cryptosporidium = 3, MNV = 2, Nippostrongylus = 2, SFB = 2, Yersinia = 3. *Candida infection includes 1 IEC replicate.
