## Supplementary Figure 6 for "Diverse infections transcriptionally reprogram the intestinal epithelium and epithelial-immune cell interactions"

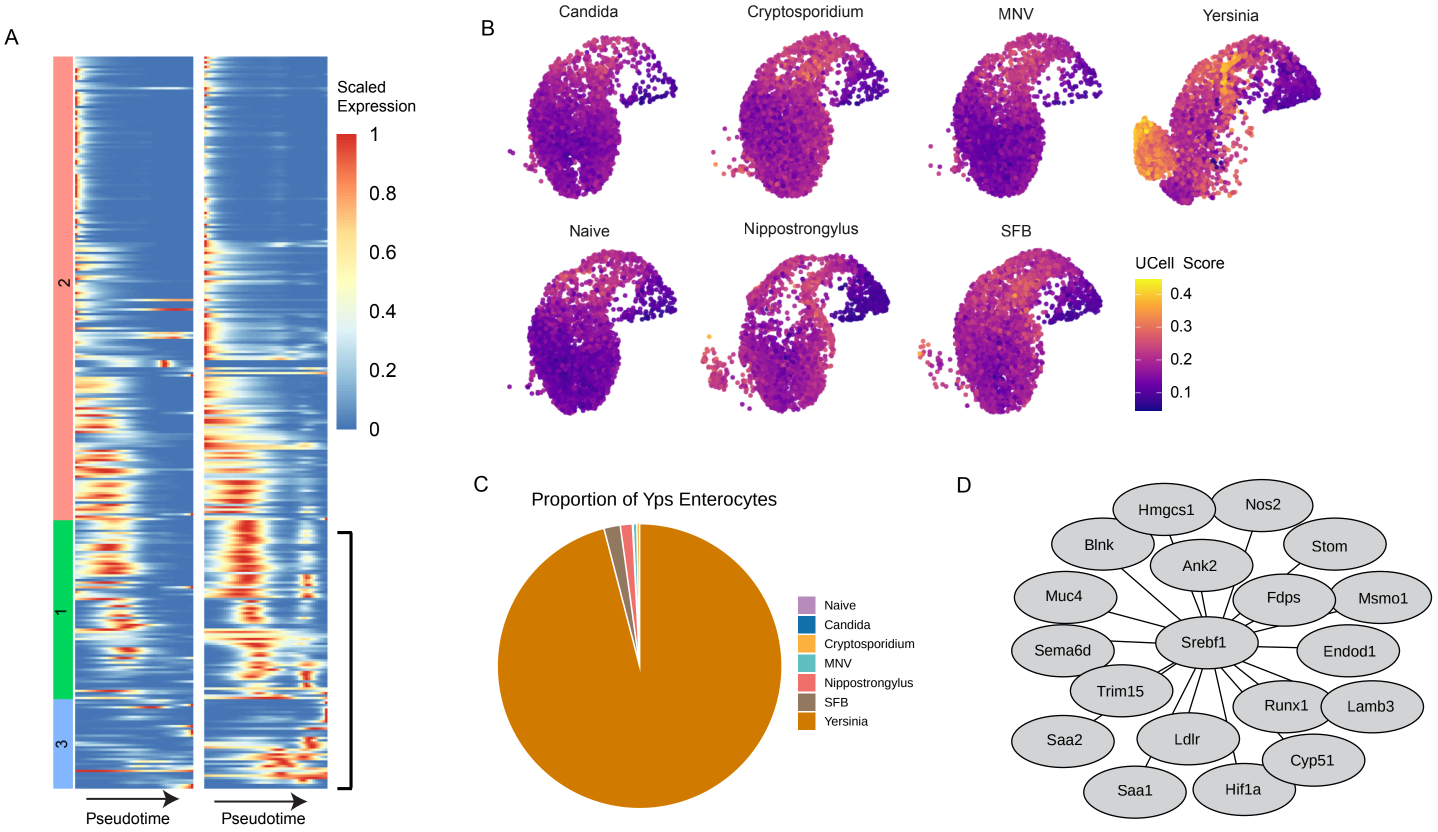


Figure S6: Unique enterocytes are found within *Yersinia*-infected samples. A) Pseudotime scaled expression of tradeSeq-defined genes that are differentially expressed along pseudotime between *Yersinia* and Naive ISC, transit amplifying, and enterocyte populations. B) Enrichment scores (color scale) (Ucell) of selected genes from Yersinia DE analysis (A, bracket) and plotted in UMAP dimensional reduced space. ISC, transit amplifying, and enterocyte populations are shown. C) Proportional frequencies of cells from each condition within the Yps enterocyte group. D) Rcistarget-defined TF network showing the motif-enriched Srebf1 TF and Yps enterocyte-defining genes containing associated Srebf1 motifs. Biological n: Naive = 2, Candida = 2*, Cryptosporidium = 3, MNV = 2, Nippostrongylus = 2, SFB = 2, Yersinia = 3. *Candida infection includes 1 IEC replicate.
