## Supplementary Figure 7 for "Diverse infections transcriptionally reprogram the intestinal epithelium and epithelial-immune cell interactions"

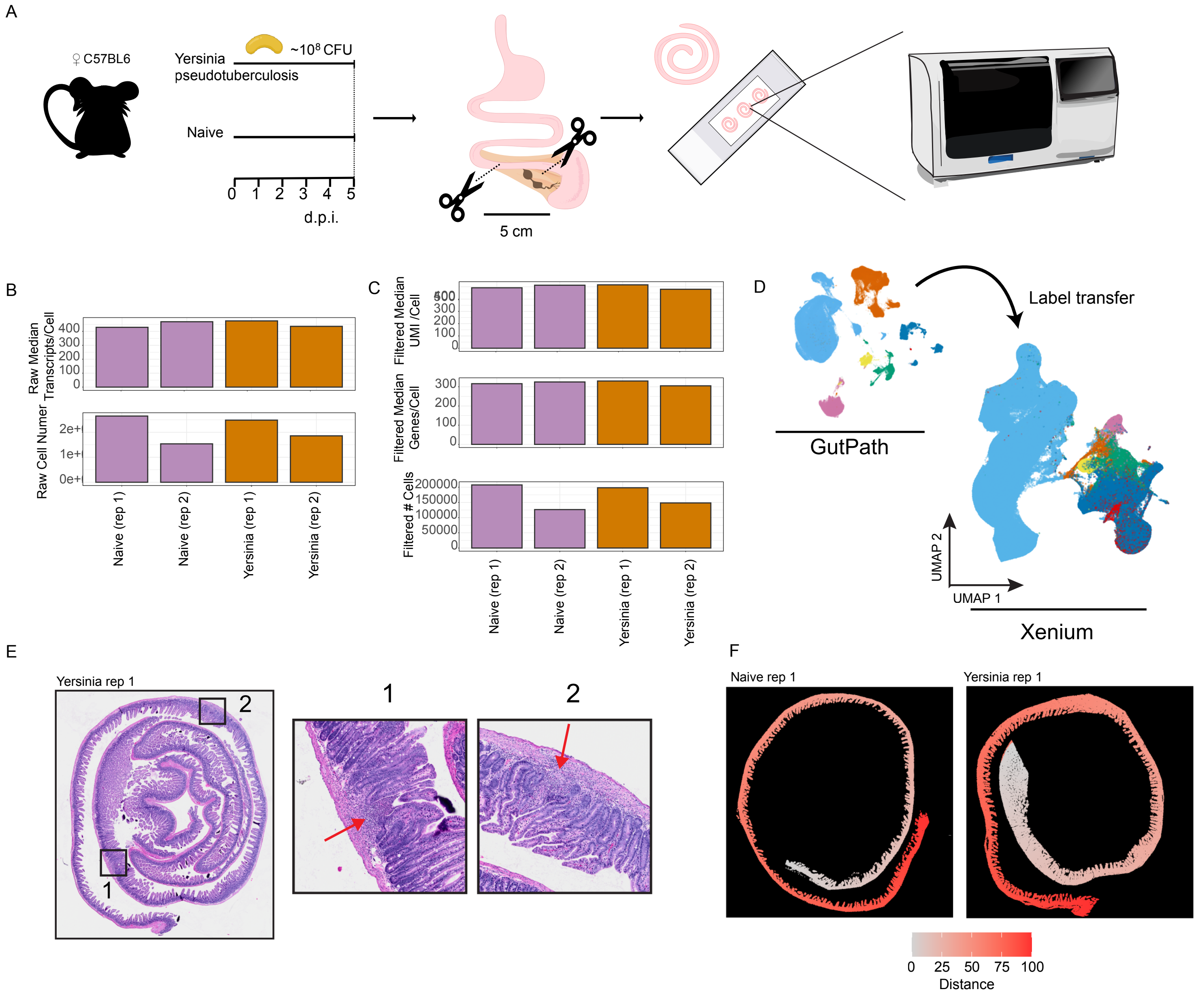


Figure S7: Pyogranuloma formation is linked to enterocyte transcriptional responses and phenotype during *Yersinia pseudotuberculosis* infection. A) Schematic diagram of *Yersinia pseudotuberculosis* infection, intestinal isolation of 5cm from the ileum, Swiss rolling, and spatially resolved transcriptomic imaging. Median raw transcripts detected per cell in each sample (top). Raw estimated cell numbers per section/sample following cell segmentation (bottom). C) Summary statistics following quality control and cell filtering: median transcripts per cell (top), median genes per cell (middle), and cell number (bottom) are shown. D) Schematic depicting UMAP reduction of CITE-seq data and major cell lineages (color) mapped onto spatially resolved Xenium UMAP dimensional reduction (bottom). E) Hematoxylin and eosin staining of an adjacent tissue section (left) from formalin fixed and paraffin embedded intestinal Swiss rolls of *Yersinia*-infected mice. Putative pyogranulomas are indicated (boxes) and shown at higher magnification (right). Immune infiltrate (red arrows) is indicated. F) Images of bioinformatically unrolled Swiss roll sections from Naive (left) and *Yersinia*-infected (right) Xenium samples. Distance (color hue) from proximal (0) to distal (100) of unrolled regions is indicated. Xenium data are representative of 2 conditions (naive, *Yersinia*) each with 2 biological replicates from different mice.
